# A Mechanically Resilient Soft Hydrogel Improves Drug Delivery for Treating Post-Traumatic Osteoarthritis in Physically Active Joints

**DOI:** 10.1101/2024.05.16.594611

**Authors:** Nitin Joshi, Jing Yan, Mickael Dang, Kai Slaughter, Yufeng Wang, Dana Wu, Trevor Ung, Virja Pandya, Mu Xian Chen, Shahdeep Kaur, Sachin Bhagchandani, Haya A Alfassam, John Joseph, Jingjing Gao, Mahima Dewani, Ryan Chak Sang Yip, Eli Weldon, Purna Shah, Chetan Shukla, Nicholas E Sherman, James N Luo, Thomas Conway, James P. Eickhoff, Luis Botelho, Ali H Alhasan, Jeffrey M Karp, Joerg Ermann

## Abstract

Intra-articular delivery of disease-modifying osteoarthritis drugs (DMOADs) is likely to be most effective in early post-traumatic osteoarthritis (PTOA) when symptoms are minimal and patients are physically active. DMOAD delivery systems therefore must withstand repeated mechanical loading without affecting the drug release kinetics. Although soft materials are preferred for DMOAD delivery, mechanical loading can compromise their structural integrity and disrupt drug release. Here, we report a mechanically resilient soft hydrogel that rapidly self-heals under conditions resembling human running while maintaining sustained release of the cathepsin-K inhibitor L-006235 used as a proof-of-concept DMOAD. Notably, this hydrogel outperformed a previously reported hydrogel designed for intra-articular drug delivery, used as a control in our study, which neither recovered nor maintained drug release under mechanical loading. Upon injection into mouse knee joints, the hydrogel showed consistent release kinetics of the encapsulated agent in both treadmill-running and non-running mice. In a mouse model of aggressive PTOA exacerbated by treadmill running, L-006235 hydrogel markedly reduced cartilage degeneration. To our knowledge, this is the first hydrogel proven to withstand human running conditions and enable sustained DMOAD delivery in physically active joints, and the first study demonstrating reduced disease progression in a severe PTOA model under rigorous physical activity, highlighting the hydrogel’s potential for PTOA treatment in active patients.

## Introduction

Post-traumatic osteoarthritis (PTOA) is the result of a primary injury to knee, ankle or other joints and represents nearly 12% of all osteoarthritis (OA)^1^. Current treatment strategies focus on symptomatic relief ^2^, and no therapeutics are available in the clinic to prevent the development or halt the progression of PTOA ^3^. Eventually, joint replacement surgery may be needed to control symptoms and restore function ^4^.

Disease-modifying osteoarthritis drugs (DMOADs) have shown promise for inhibiting or reversing PTOA progression in pre-clinical studies ^5^. However, the clinical translation of DMOADs has been lagging behind; one reason being their short half-live in joints following intra-articular administration^6^. This necessitates their systemic administration which is associated with an increased risk for off-target side effects. The half-life of locally injected therapeutics in joints can be increased with intra-articular drug delivery systems that exhibit sustained drug release. However, strenuous activities such as running or playing sports can be detrimental for these delivery systems^7^ causing pre-mature drug release during physical activity and/or faster release afterwards, thereby reducing the overall joint residence time and efficacy of the drug. This is particularly relevant for DMOADs, as they are likely to work best in early disease^8^, when symptoms are minimal and patients are physically active. Traumatic injuries often occur in young adults with symptoms of PTOA beginning within 5 years after the injury and documentation of early cartilage degeneration in the majority of these individuals in their 30s and 40s^9^. Although previously developed sustained release platforms, including hydrogels ^10–14^, nano- and micro-particles ^15–19^, have shown pre-clinical promise for intra-articular delivery of DMOADs, none have been evaluated in physically active joints nor have they considered the impact of mechanical loading on drug release.

Mechanically tough materials, such as polyethylene and polyetheretherketone can withstand mechanical loading in joints ^20,21^. However, sustained wear of these materials over time due to mechanical friction results in the formation of debris and small particles which may elicit an inflammatory response ^22^. Soft materials may be better suited for DMOAD delivery as they can also provide a lubricating effect in joints ^23^. However, mechanical loading can cause architectural damage in soft materials ^24^, which may negatively impact DMOAD release.

In this study, we developed a soft hydrogel which exhibits mechanical resilience, thereby minimizing the impact of mechanical loading on the release kinetics of the DMOAD, and enabling efficient DMOAD delivery under conditions of physical activity. We used triglycerol monostearate (TG-18), a low molecular weight gelator (LMWG), which self-assembles to form a soft hydrogel. Due to its supramolecular nature, TG-18 hydrogel shows dynamic self-healing, known as thixotropy, which enables mechanical resilience. Moreover, compared to supramolecular polymeric hydrogels, LMWG -based hydrogels such as TG-18 exhibit faster self-healing owing to greater molecular mobility of LMWG compared to polymers^25^, enabling rapid recovery. During mechanical loading, the TG-18 hydrogel transitions from a gel to a viscous fluid state, but rapidly reverts to the gel state upon removal of the mechanical load, thereby regaining its original viscoelastic and morphological properties.

TG-18 hydrogel stably encapsulates a wide range of therapeutics releasing them in a sustained manner due to gradual hydrolysis of the ester bond in TG-18 ^26–30^. For this proof-of-concept study, we encapsulated the cathepsin-K inhibitor L-006235 non-covalently into TG-18 hydrogel (Fig. 1) because of its previously reported efficacy in pre-clinical OA models upon oral administration, and established mechanism of action^31–36^. We show that L-006235-loaded TG-18 hydrogel, hereafter referred to as L-006235 gel, showed rapid recovery following mechanical loading conditions that simulate human running. It showed no premature release of L-006235 during mechanical loading and no impact on the release kinetics of L-006235 afterwards. In contrast, a soft polymeric hydrogel reported previously for intra-articular drug delivery^37,38^ and used as a control in our study failed to recover, and showed increased drug release following mechanical loading.

**Fig. 1.**
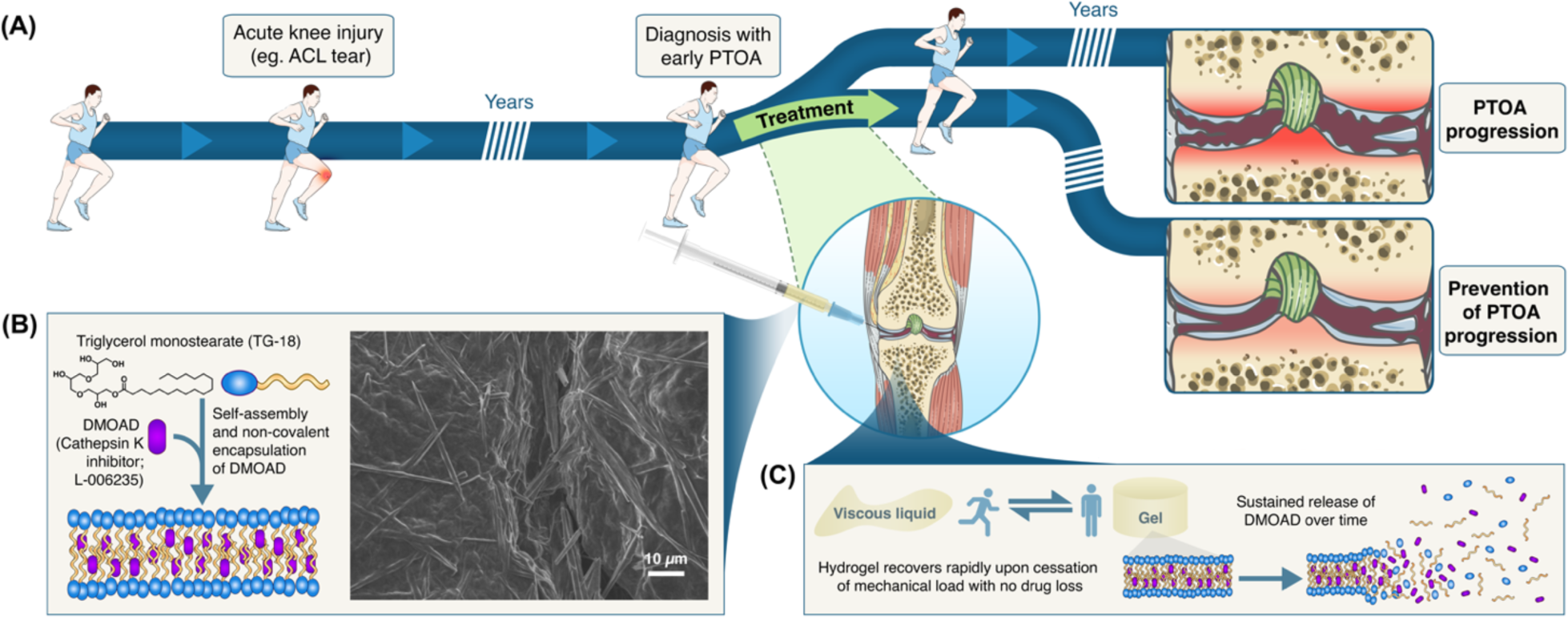
DMOADs can be encapsulated into triglycerol monostearate (TG-18) hydrogel for the prevention of PTOA progression in active joints. **(A)** Schematic depicting intra-articular delivery of DMOAD-loaded TG-18 hydrogel as a strategy to prevent disease progression in young, active patients diagnosed with early PTOA. **(B)** The cathepsin-K inhibitor L-006235 was used as a proof of concept, hydrophobic DMOAD in this study. TG-18 self-assembles into lamellar structures and encapsulates L-006235 *via* non-covalent, hydrophobic interactions. High-resolution scanning electron microscopy (HR-SEM) image of L-006235 gel demonstrating higher order fibrous assemblies that form the bulk hydrogel. **(C)** Schematic showing sustained release of the DMOAD from TG-18 hydrogel due to diffusion and gradual hydrolysis of TG-18. The viscoelastic properties of the intraarticular hydrogel recover rapidly after mechanical loading, with no pre-mature release or loss of L-006235 during mechanical loading, and no impact on the release kinetics afterwards. Parts of the figure were drawn by using pictures from Servier Medical Art. Servier Medical Art by Servier is licensed under a Creative Commons Attribution 3.0 Unported License (https://creativecommons.org/licenses/by/3.0/).

Notably, TG-18 hydrogel also demonstrated robust lubricating properties under varied mechanical loading conditions and reduced the coefficient of friction between cartilage discs, as confirmed *in vitro*. Hydrogel injected into mouse knee joints showed similar release kinetics of a hydrophobic fluorescent dye in treadmill-running *versus* non-running mice and sustained release of L-006235 was observed in treadmill-running mice. Using a mouse model of aggressive PTOA, with disease exacerbated by rigorous treadmill running we show that L-006235 gel prevented disease progression more effectively than free L-006235. Together, our findings underscore the potential of TG-18 hydrogel as a mechanically resilient platform for enabling effective intra-articular delivery of DMOADs to prevent the progression of PTOA in physically active patients. To our knowledge, this is the first hydrogel proven to withstand human running conditions and enable sustained delivery of a DMOAD in physically active joints. It is also the first study to demonstrate reduced disease progression in an aggressive PTOA model exacerbated by rigorous physical activity.

## Results

### TG-18 hydrogel self-assembles to encapsulate L-006235 and exhibits sustained drug release

TG-18 is a LMWG with a hydrophilic polyhydroxyl head group and a hydrophobic polymethylene tail group (Fig. 1B). Heating TG-18 to a temperature of 55-60°C or higher in a DMSO/water mixture results in its dissolution by micelle formation ^26^. Hydrophobic drugs such as L-006235 can be added at this time. Cooling the drug-LMWG mixture results in self-assembly of TG-18 into higher order fibrous structures that entangle to form the bulk gel ^26^. During the self-assembly process, L-006235 is incorporated into the core of the fibers *via* hydrophobic interactions between L-006235 and the stearic acid chains of TG-18. We studied the hydrophobic interactions between TG-18 and L-006235 using *in silico* molecular docking. This analysis yielded nine possible conformers with ΔG varying from −3.4 to −3 kCal/mol (Table S1), negative ΔG values indicating spontaneous interaction between TG-18 and L-006235. The 3D visualization of the molecular docking showed Pi-Sigma hydrophobic interactions between the hydrocarbon chain of TG-18 and the aromatic ring of L-006235 with a distance of 3.84 Å (Fig. S1A). Van der Waals interactions were also observed between L-006235 and TG-18 (Fig. S1B).

We prepared TG-18 hydrogel (10% w/v) and encapsulated L-006235 at a final concentration of 10, 15 or 20 mg/mL. Hydrogel containing 10 or 15 mg/mL of L-006235 showed fibrous morphology similar to blank hydrogel when analyzed with high-resolution scanning electron microscopy (HR-SEM) (Fig. 1B, Fig. S2, A-C). At 20 mg/mL of L-006235, non-fibrous precipitates were observed (Fig. S2D) indicating unsuccessful gelation. Therefore, we chose 15 mg/mL as the optimal loading concentration of L-006235 for all subsequent *in vitro* and *in vivo* studies.

L-006235 gel was easily injectable through a 27 gauge needle (Movie S1), which we used for intra-articular injection in mouse joints. Since intra-articular injection in larger joints are performed using larger needles (21 gauge) ^39^, we do not anticipate any injectability issues in humans knee joints. We evaluated the release of L-006235 from L-006235-gel upon exposure to synovial fluid from human OA joints *in vitro*. We observed sustained release of L-006235, with similar release kinetics in phosphate buffered saline (PBS) (pH 7.4) without or with addition of OA synovial fluid (Fig. S3). Less than 45% cumulative release of L-006235 was observed over a period of 30 days.

### L-006235 gel recovers rapidly following mechanical loading at levels relevant to running human knees, with no impact on drug release

Using a rotational rheometer, we performed strain sweep and dynamic frequency sweep measurements to characterize the viscoelastic properties and define the linear viscoelastic (LVE) region for blank TG-18 hydrogel and L-006235 gel ^40^. The LVE region indicates the range of strain that can be applied to a material without impacting its physical integrity. Within the LVE region, a hydrogel maintains its gel-like state and the shear storage modulus (G’) exceeds the shear loss modulus (G”), i.e. G’>G”. The storage modulus represents the energy stored in the elastic part of a material and indicates the amount of structure present in it. The loss modulus represents the viscous part or the amount of energy dissipated during deformation. We varied the strain from 0.01% to 50%, while keeping the frequency constant at 0.1 Hz, in order to determine the flow point of the hydrogel, i.e. the point where the hydrogel transitions from gel to a viscous liquid, resulting in G’<G”. L-006235 gel showed a lower flow point compared to the blank hydrogel (5% *versus* 22% strain) (Fig. S4, A and C). However, L-006235 gel exhibited 10-fold higher values of G’ and G” in the LVE region compared to the blank hydrogel, suggesting improved mechanical strength due to drug loading. Next, a dynamic frequency sweep was performed from 0.1 to 5 Hz, while keeping the strain constant at 0.1%, which was within the LVE region of both the blank and L-006235-loaded hydrogel. G′ remained higher than G” over the entire frequency sweep and an increase in modulus was observed with increasing frequency (Fig. S4, B and D).

We then evaluated the impact of repeated mechanical loading on the viscoelastic properties of L-006235 gel and *in vitro* release of L-006235 using experimental conditions that model human knees under resting and running conditions ^41,42^. L-006235 gel was subjected to low strain and low frequency (0.5%, 0.1 Hz; conditions resembling a resting human knee) for 1 minute followed by 30 consecutive cycles of alternating strain and frequency, during which G’ and G” were measured. Each cycle involved 1 minute of high strain and high frequency (35%, 2.5 Hz; conditions resembling a running human knee) followed by 1 minute of low strain and low frequency (Fig. 2A). Our experiment therefore simulated 60 minute of interval running by humans, involving 1 min of running at the rate of 150 steps/min, followed by 1 min of resting, repeated over 30 cycles. Under low strain and low frequency conditions, the hydrogel maintained its gel-like state with G’>G” (Fig. 2B). This is expected as the low strain and low frequency are within the LVE region of the L-006235 gel. Application of high strain and high frequency resulted in loss of the gel-like state and transition of the hydrogel to a viscous liquid state, with G’<G”. Upon removal of high strain and high frequency, the hydrogel rapidly reverted to the gel-like state and regained its original viscoelastic properties (Fig. 2B). This thixotropic behavior was observed regardless of the number of cycles applied (up to 30 cycles). Release of L-006235 from the hydrogel following repeated cycles of alternating strain and frequency was largely indistinguishable from that of a fresh hydrogel (Fig. 2C), indicating that mechanical loading had no impact on L-006235 release kinetics. L-006235 gel subjected to mechanical loading and later centrifuged to separate free drug, was found to have a similar concentration of L-006235 as fresh L-006235 gel (Fig. S5). This indicates that high strain and high frequency conditions did not result in pre-mature release and loss of L-006235 from hydrogel during its viscous state. Finally, HR-SEM imaging performed following the application of mechanical loading revealed maintenance of the fibrous morphology (Fig. 2D).

**Fig. 2.**
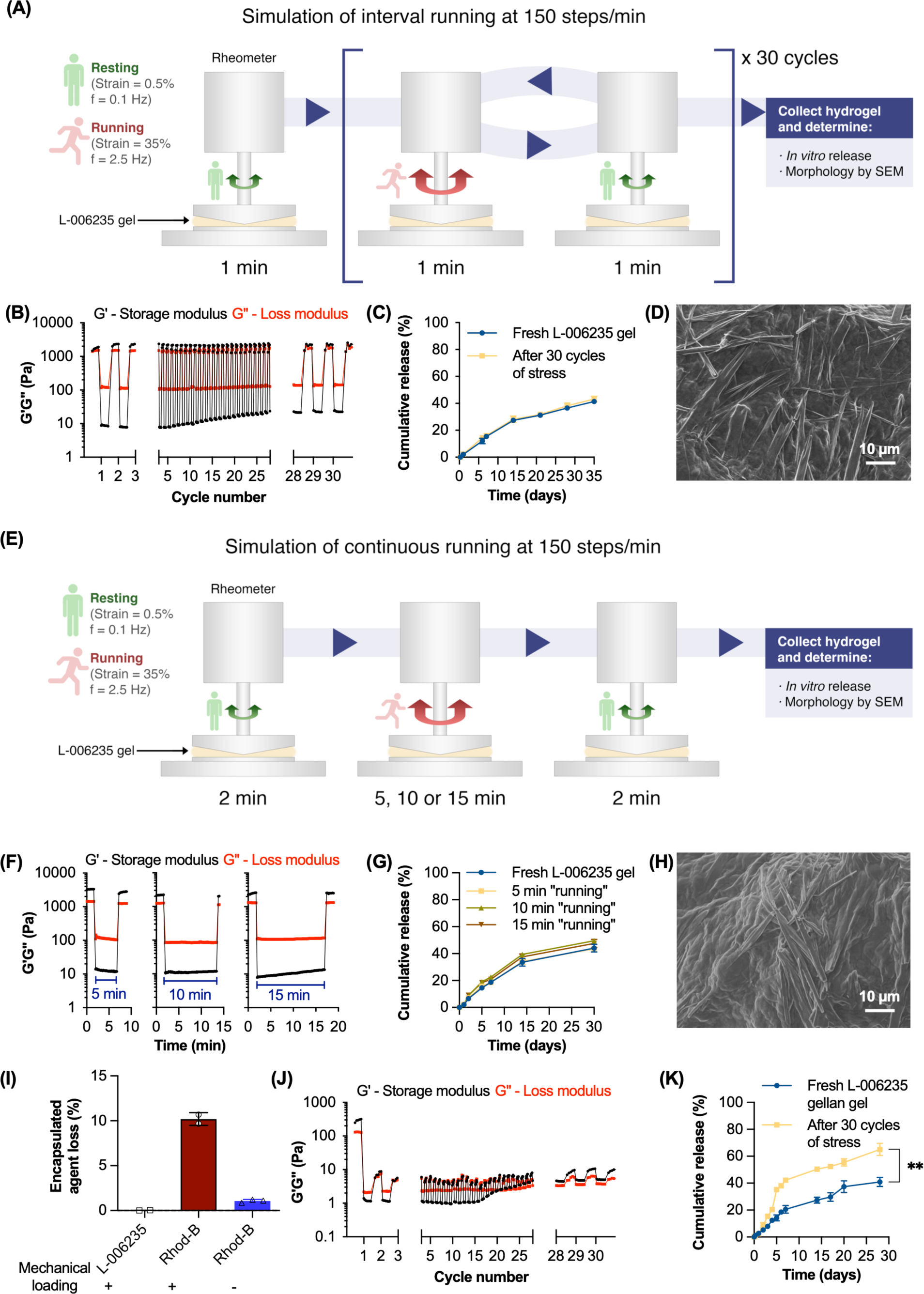
L-006235 gel recovers rapidly following mechanical loading conditions simulating human running. **(A)** Experimental outline to measure the effect of repeated mechanical loading conditions simulating interval running by humans on L-006235 gel, and release kinetics of L-006235. Using a rotational rheometer, the L-006235 gel was subjected to 30 cycles of alternating strain and frequency in the presence of PBS. Each cycle involved 1 minute of high strain and high frequency relevant to a running human knee followed by 1 minute of low strain and low frequency relevant to a resting human knee. **(B)** Shear storage modulus (G’) and shear loss modulus (G”) were recorded. **(C and D)** After 30 cycles, L-006235 gel was collected, evaluated for *in vitro* drug release in PBS (37°C) and analyzed for morphology by HR-SEM. **(E)** Experimental outline to determine the effect of a single cycle of alternating strain and frequency on L-006235 gel at 37°C in the presence of PBS. The cycle involved 2 minutes of low strain and low frequency followed by high strain and high frequency for 5, 10 or 15 minutes, and finally low strain and low frequency for 2 minutes. **(F)** Shear storage modulus (G’) and shear loss modulus (G”) were measured. **(G)** At the end of each cycle, L-006235 gel was collected and evaluated for *in vitro* drug release in PBS (37°C)**. (H)** Gel morphology was assessed using HR-SEM after a loading cycle with a 15 min step of high strain and high frequency. **(I)** Fraction of L-006235 or rhodamine-B (Rhod-B) released from TG-18 hydrogel in the presence of PBS during a single cycle of mechanical loading with a 15 min step of high strain and high frequency, and fraction of Rhod-B released in the absence of mechanical loading. **(J)** Shear storage modulus (G’) and shear loss modulus (G”) of L-006235-loaded gellan gel (L-006235 gellan gel) measured during 30 consecutive cycles of alternating strain and frequency at 37°C. **(K)** After 30 cycles, L-006235 gellan gel was collected and evaluated for *in vitro* drug release in PBS at 37°C (\*\**P* < 0.01 on day 28). Data in **C, G, I, and K** are means ± SD of technical repeats (n = 3, each experiment performed at least twice). *P* value in **K** was determined using Student’s *t*-test. Data in **B, F, and J** are from a single experiment (experiment repeated three times).

L-006235 gel also showed rapid recovery following high strain and high frequency conditions simulating continuous running by humans at 150 steps/min for 5, 10, or 15 min (Fig. 2, E and F), with no changes in the drug release kinetics or in the fibrous morphology as compared to a fresh hydrogel (Fig. 2, G and H). Gel collected after each cycle had the same concentration of L-006235 as fresh L-006235 gel, indicating that no pre-mature release and loss of L-006235 occurred during the viscous state of the hydrogel (Fig. S6). In a control experiment, we encapsulated rhodamine-B, a hydrophilic dye into TG-18 hydrogel, and subjected the rhodamine-B-loaded hydrogel (Rhod-B gel) to a single 15 minute cycle of high strain and high frequency. While Rhod-B gel showed thixotropic properties similar to L-006235 gel (Fig. S7), we observed ∼10% release of rhodamine-B from Rhod-B gel under mechanical loading conditions (Fig. 2I). This is consistent with the inability of hydrophilic rhodamine-B to interact and bind to TG-18 leading to pre-mature release during the viscous state. Compared to Rhod-B gel under mechanical loading, significantly less release of rhodamine-B was observed in the absence of mechanical loading (Fig. 2I), confirming that the pre-mature loss of rhodamine-B under mechanical loading conditions is not simply due to the release of the dye present at the surface of the hydrogel.

Similar to L-006235 gel, blank hydrogel also showed rapid recovery of viscoelastic properties following high strain and high frequency conditions (Fig. S8), suggesting that TG-18 hydrogel is inherently thixotropic and can rapidly recover from levels of repeated mechanical loading relevant to the running human knee. Rheo-microscopy demonstrated fibrous or thread like structures in L-006235 gel under both low strain and low frequency, and high strain and high frequency conditions (Fig. S9), providing direct, visual confirmation that even in the viscous state under high strain and high frequency conditions, TG-18 hydrogel maintains its higher order fibrous structure.

We then demonstrated that the thixotropic behavior of TG-18 hydrogel is the result of self-assembly of TG-18 into a supramolecular hydrogel, and not an inherent property of the TG-18 molecule. To this end, TG-18 was dissolved in DMSO (TG-18/DMSO; 10% w/v) rather than DMSO/water by heating the mixture to 60-80°C, followed by cooling for 15-30 min at room temperature. This process results in a non-assembled TG-18/DMSO mixture, instead of a self-assembled hydrogel. Unlike TG-18 hydrogel, the TG-18/DMSO mixture did not fully recover its viscoelastic properties upon removing high strain and frequency (Fig. S10). The storage modulus kept decreasing with each consecutive cycle of alternating strain and frequency. This provides evidence that self-assembly of TG-18 into an organized supramolecular hydrogel is critical to achieve the thixotropic behavior.

We next encapsulated L-006235 in a gellan gum-based soft hydrogel reported previously for intra-articular drug delivery^37,38^. Unlike L-006235 gel, L-006235 gellan gel failed to recover its gel-like state upon removing high strain and frequency, which is consistent with its non-thixotropic nature (Fig. 2J). The mechanical integrity of the gel was lost during the first cycle, with a 100-fold reduction in both the storage and loss modulus. Moreover, L-006235 release from gellan hydrogel exposed to repeated cycles of alternating strain and frequency was significantly higher than from a fresh gellan hydrogel (Fig. 2K). Our data demonstrate the superiority of the TG-18 hydrogel over the previously developed gellan gum-based hydrogel for drug delivery under mechanical loading conditions. Additionally, our findings establish that thixotropy is essential for maintaining the release kinetics of L-006235 from a hydrogel when subjected to mechanical loading.

### L-006235 gel is biocompatible with cells from human joints and safe for intra-articular administration in mice

As a prelude to *in vivo* studies, we investigated the *in vitro* biocompatibility of L-006235 gel with primary human chondrocytes and fibroblast-like synoviocytes from healthy donors and OA patients. Cells were incubated for 24, 48, or 72 h in medium, medium containing L-006235 gel or free L-006235 at a 2 nM final concentration of L-006235, which is 8 times higher than the IC_50_ of L-006235 (0.25 nM, information provided by the manufacturer). Cells incubated in PBS, medium with blank hydrogel or DMSO/water (20% v/v DMSO) added at volumes equivalent to L-006235 gel were evaluated as controls. As expected, cells incubated in PBS exhibited a significant reduction in metabolic activity compared to cells incubated in medium (Fig. 3, A-D). Incubation in medium containing L-006235 gel, blank hydrogel, DMSO/water or free L-006235 resulted in a maintenance of 80% or greater metabolic activity relative to cells incubated in culture medium alone at all time points. We also evaluated the biocompatibility of hydrogel formulations at a 200 nM and 2 µM final concentration of L-006235, which is 800 and 8000-fold higher than its IC_50,_ respectively. Even under those conditions, close to 100% metabolic activity was observed for L-006235 gel in all four cell types (Fig. S11, A-D), which confirms its biocompatibility based on published reports for assessing *in vitro* toxicity ^43^. Although we observed only 75% metabolic activity for free L-006235 at 2 µM concentration, improved biocompatibility of L-006235 in the hydrogel formulation is most likely due to the sustained release of L-006235, and further emphasizes the advantage of our hydrogel system for sustained delivery of DMOADs.

**Fig. 3.**
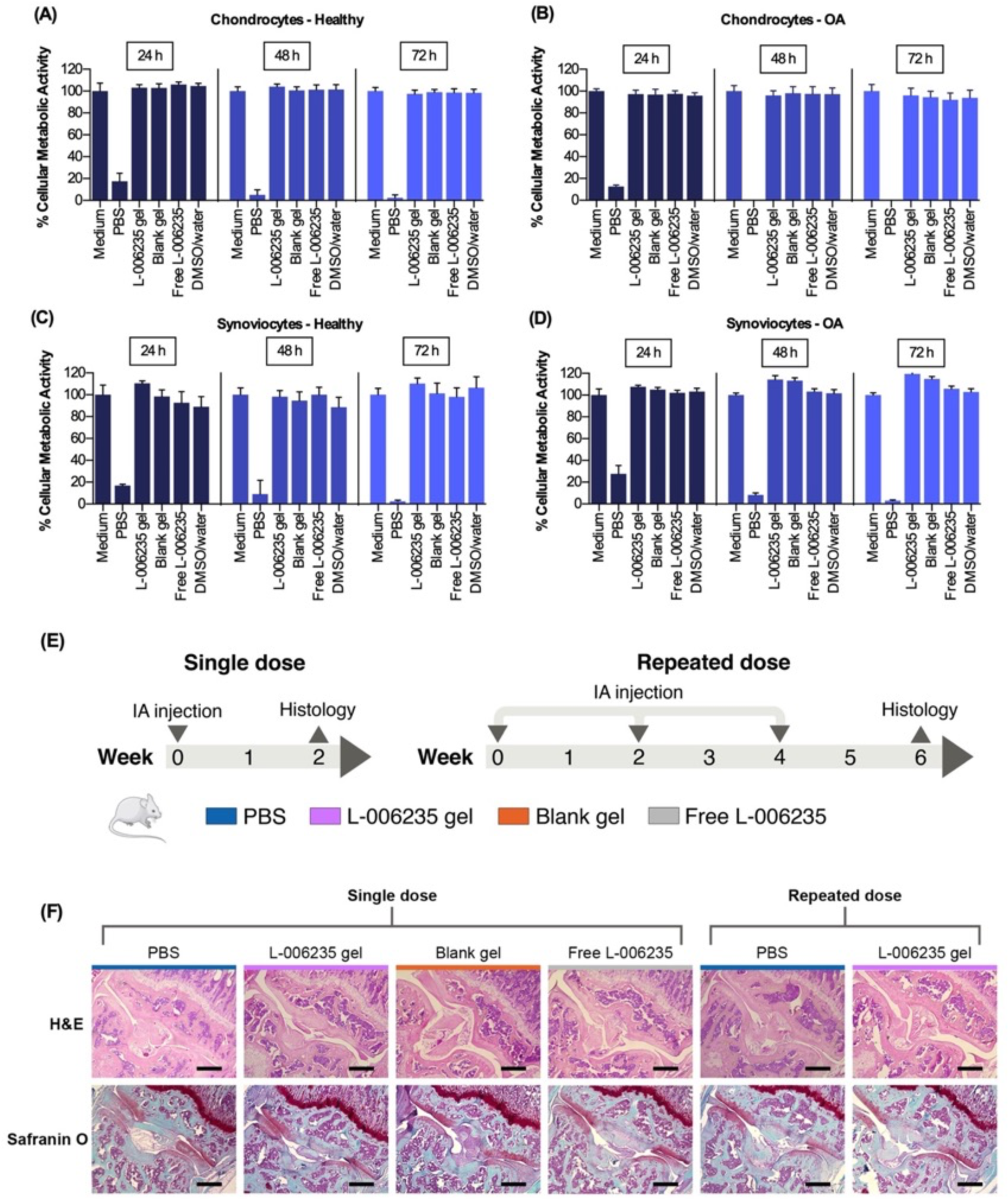
L-006235 gel is biocompatible with cells from human joints and safe for single and repeated intra-articular administration in mice. **(A-D)** Primary human chondrocytes and fibroblast-like synoviocytes from healthy donors or OA patients were incubated in a 96-well plate in medium, PBS, or in medium with L-006235 gel or free L-006235 at a 2 nM final concentration of L-006235. Cells incubated in medium with blank TG-18 hydrogel (Blank gel) or DMSO/water at volumes equivalent to L-006235 gel were included as controls. After 24, 48 or 72 h of incubation, metabolic activity was determined by XTT assay. **(E)** Experimental outline to evaluate the safety of single or repeated intra-articular injections of L-006235 gel (4 µl, 60 µg L-006235) in mice. In the single dose study, healthy mice were injected on day 0 with 4 µL PBS, L-006235 gel, blank gel or free L-006235. Two weeks later, injected knees were harvested and processed for histology. In the repeated dose study, knees were injected with 4 µl PBS or L-006235 gel on day 0, week 2 and week 4, and harvested for histology at week 6. **(F)** Representative images of histological sections stained with H&E or safranin O from each experimental group (scale bar: 200 µm). Data in **A-D** are means ± SD of technical repeats (n = 8 wells per condition, experiment performed at least twice).

Next, we assessed the *in vivo* safety of L-006235 gel in healthy C57BL/6J mice injected intraarticularly either once or repeatedly (Fig. 3E). Analysis of knee sections was performed using Hematoxylin and eosin (H&E) and Safranin O staining, which have been previously used as standard techniques to assess biocompatibility in knee joints ^44,45^. H&E stained sections of knees from all groups in both the single-dose and repeated-dose studies did not show any inflammation or other gross evidence of toxicity (Fig. 3F). Safranin O staining also did not reveal any detrimental effects on the cartilage (Fig. 3F).

### Release of encapsulated agents from hydrogel injected into mouse joints is not affected by treadmill running

Having demonstrated that both blank and L-006235-loaded hydrogel can rapidly recover their viscoelastic properties following mechanical loading *in vitro*, we investigated the impact of mechanical loading on the release of encapsulated agents from TG-18 hydrogel *in vivo.* To this end, hydrogel was co-loaded with L-006235 and the hydrophobic fluorescent dye 1,1′-dioctadecyl-3,3,3′,3′-tetramethylindotricarbocyanine iodide (DiR) (4 µL, 60 µg L-006235, 0.4 µg DiR) DiR signals can be easily monitored in live mice using an *in vivo* imaging system (IVIS). *In vitro*, DiR showed similar release kinetics as L-006235 from the co-loaded hydrogel (Fig. S12) confirming the suitability of DiR as a surrogate marker for the *in vivo* monitoring of drug release. Hydrogel co-loaded with L-006235 and DIR was then injected into knee joints of C57BL/6 mice. One group of mice ran on a treadmill at 400 m/30 min/day ^46^, 5 days/week while the other group did not run (Fig. 4A). Mice were imaged every other day using IVIS. Both groups demonstrated similar release kinetics of DiR – measured as the loss of fluorescence signal over time (Fig. 4, B-D). These results illustrate that mechanical loading induced by treadmill running does not affect the release of encapsulated agents from the hydrogel.

**Fig. 4.**
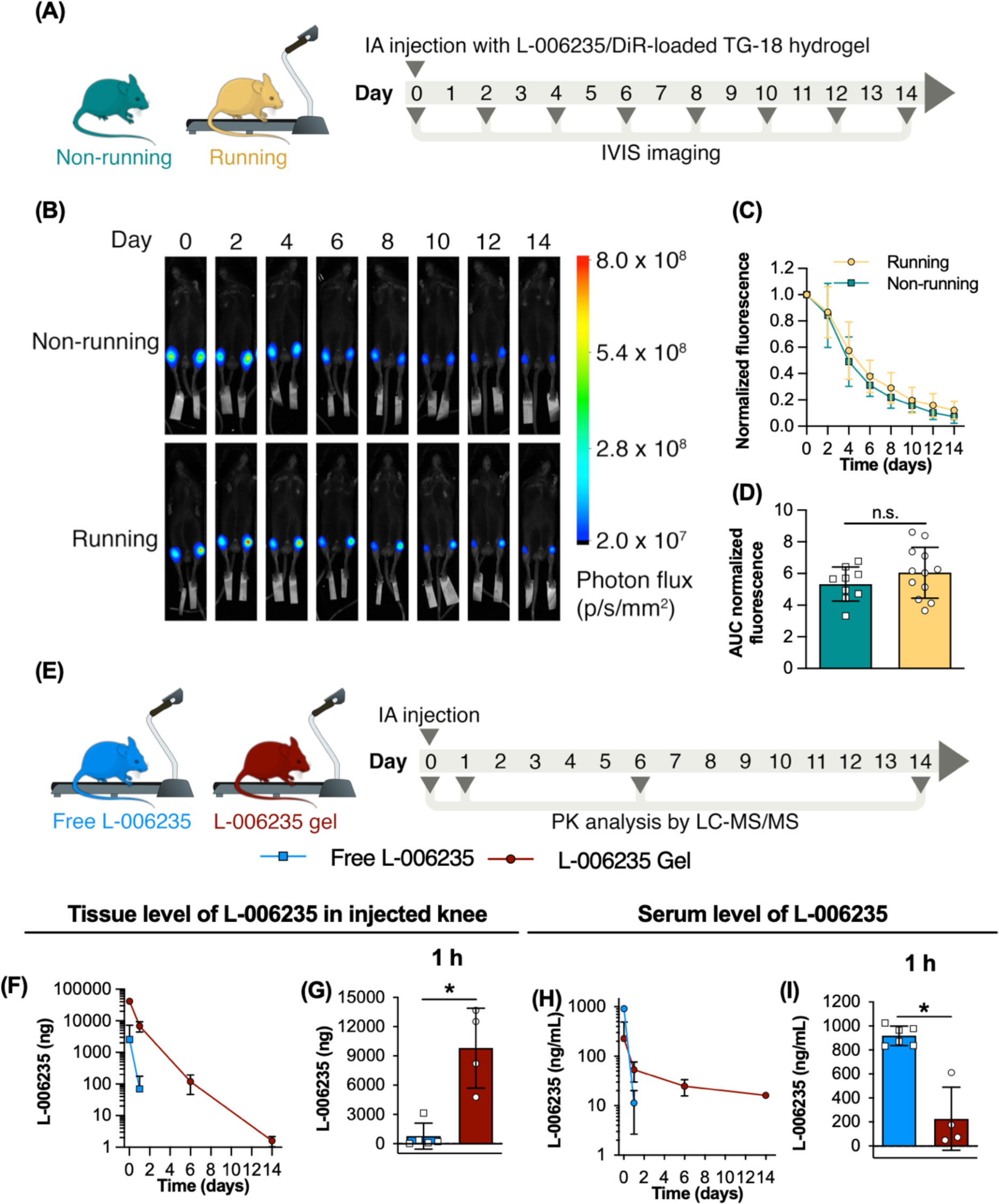
Treadmill running does not impact release kinetics of encapsulated agents from TG-18 hydrogel injected into mouse knee joints. **(A)** Experimental outline to assess the impact of treadmill running on release of an encapsulated hydrophobic fluorescent dye (DiR) from TG-18 hydrogel. On day 0, mice were injected into both knees with TG-18 hydrogel co-loaded with DiR and L-006235 (4 µl, 60 µg L-006235 and 0.4 µg DiR). Subsequently, one group ran on a treadmill at 400 m/30 min/day, 5 days/week, while the other group did not run. Mice were imaged every other day using an *in vivo* imaging system (IVIS). **(B)** IVIS images of one representative animal from each experimental group. **(C and D)** Fluorescence signal (normalized to day 0) measured over the injected knees and area under the curves (AUCs). **(E)** Experimental outline of the pharmacokinetic study. L-006235 gel or free L-006235 (4 µl, 60 µg L-006235) was injected into the right knee of healthy mice. Mice subsequently ran on a treadmill at 400 m/30 min/day, 5 days/week. L-006235 level was quantified in the injected knee and in serum at 1 h, day 1, 6 and 14 post injection. **(F and G)** L-006235 level in the injected knee over time and at 1 h after injection. In mice injected with free L-006235, no drug was detected in the tissue beyond day 1 (\**P* < 0.05). **(H and I)** L-006235 serum level over time and 1 h after intra-articular injection (\**P* < 0.05). In mice injected with free L-006235, only two animals showed detectable drug levels on day 1, and subsequently no drug was detected. In mice injected with L-006235 gel, only 2 animals showed detectable drug levels on day 14. Data in **C and D** are means ± SD (n = 4-6 mice/group, experiment performed twice). *P* values were determined using Student’s *t*-test. Data in **F-I** are means ± SD (n = 3-6 mice/group, experiment performed twice). *P* values were determined using Student’s *t*-test.

To investigate the *in vivo* release kinetics of L-006235 from TG-18 hydrogel in treadmill running mice, free L-006235 or L-006235 gel (4 µL, 60 µg L-006235) was injected into the right knee joint of healthy C57BL/6J mice on day 0 (Fig. 4E). Mice then ran on a treadmill at 400 m/30 min/day, 5 days/week, and euthanized at 1 h, day 1, 6 or 14, and the injected knee and serum were collected to analyze L-006235 levels by LC-MS/MS. Mice injected with L-006235 gel showed a gradual decline of the L-006235 level in the injected knee tissue. Drug was detectable at all time points including day 14, both in the injected knee and in the serum (Fig. 4 F and H), consistent with sustained release. In mice injected with free L-006235, the tissue drug level in the injected knee declined rapidly, and no drug was detectable beyond day 1 (Fig. 4F). Even at 1 h post-injection, mice who had received free L-006235 showed a 13-fold lower drug level in the injected knee compared to mice injected with L-006235-loaded hydrogel (Fig. 4G) while serum levels of L-006235 were 5-fold higher (Fig. 4I). Together, these results demonstrate that TG-18 hydrogel results in sustained release of L-006235 in treadmill running mice and prolongs the joint residence time of the drug compared to the injection of free L-006235.

### L-006235 gel reduces PTOA progression in treadmill-running mice

We then investigated the therapeutic efficacy of L-006235 gel in treadmill running mice induced to develop PTOA through destabilization of the medial meniscus (DMM) surgery ^47^. DMM or sham surgery was performed in adult C57BL/6J mice on day 0. Three weeks later, after the surgical incision had completely healed, mice with DMM surgery were injected in the operated right knee with L-006235 gel (4 µL, 60 µg L-006235), free L-006235 (4 µL, 60 µg L-006235), blank hydrogel (4 µL) or DMSO/water (20% v/v DMSO, 4 µL), followed by repeat dosing at week 5 and 7, i.e. once every two weeks (Fig. 5A). From week 3 to week 8, all mice ran on a treadmill at 400 m/30 min/day, 5 days/week. The running speed of 400 m/30 min/day was based on a previously published report ^46^, and was chosen to exacerbate the OA phenotype in mice after DMM surgery, resulting in aggressive disease progression. Mice were euthanized at week 9 and knees were harvested. After micro-CT imaging, knees were decalcified, sectioned, and stained with Safranin O. Sections were scored for cartilage degeneration using the Osteoarthritis Research Society International (OARSI) system in a blinded manner ^48^.

**Fig. 5.**
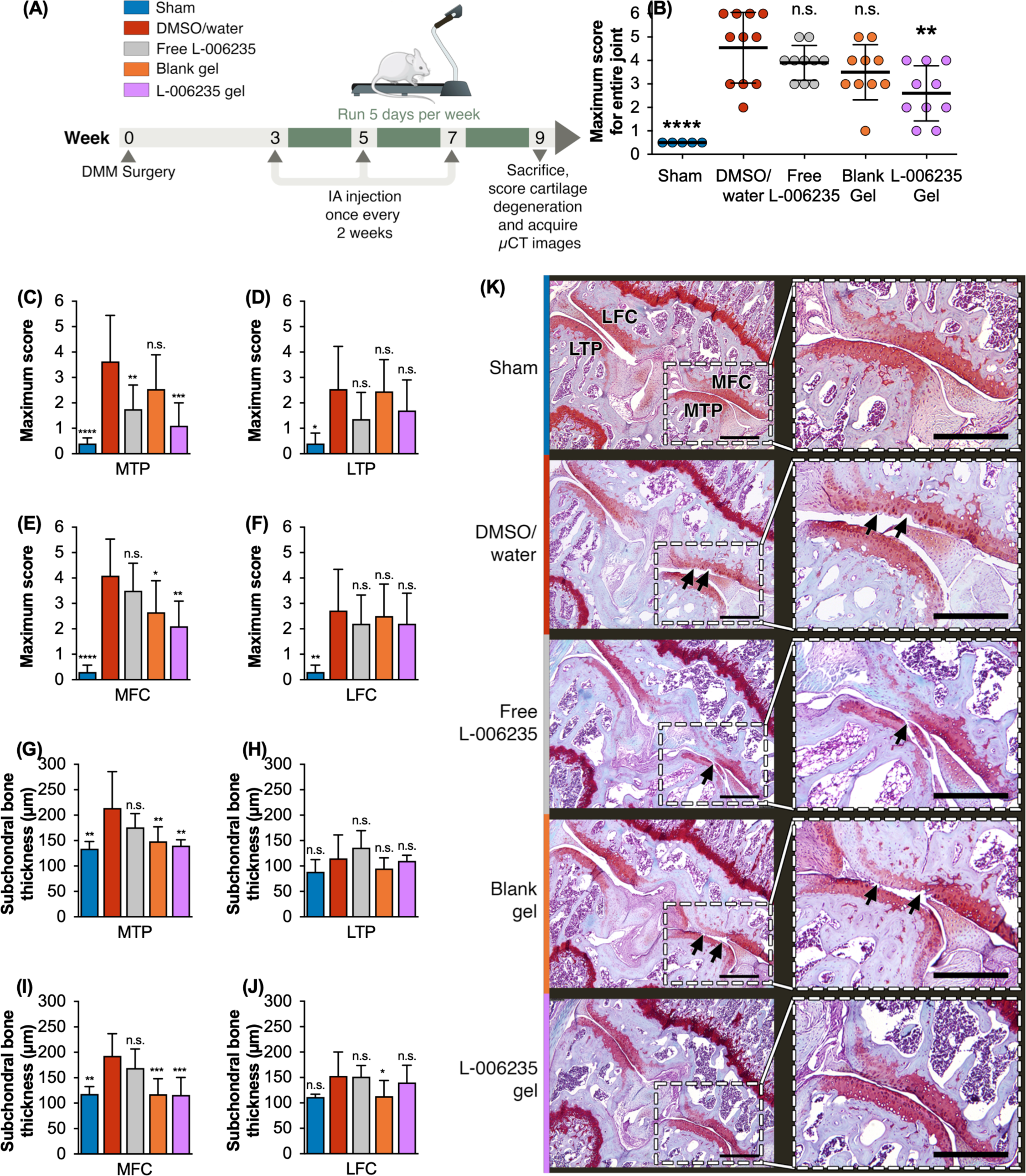
Intra-articular delivery of L-006235 in TG-18 hydrogel reduces OA severity in treadmill-running mice. **(A)** Experimental outline to determine the therapeutic efficacy of L-006235 gel in treadmill-running mice with surgically induced PTOA. Destabilization of the medial meniscus (DMM) surgery or sham surgery was performed on day 0. Three weeks later, mice with DMM surgery were injected in the right knee with L-006235 gel, free L-006235, blank TG-18 hydrogel (Blank gel, 4 µl) or DMSO/water followed by repeat dosing at week 5 and 7. From week 3 to week 8, mice from all groups ran on a treadmill at 400 m/30 min/day, 5 days/week. Mice were euthanized at week 9 and knees harvested for analysis by µCT and histology. **(B-F)** OARSI maximum scores for entire knee joints and for each quadrant of the joint – medial and lateral tibial plateau (MTP and LTP), medial and lateral femoral condyle (MFC and LFC) (\**P* < 0.05, \*\**P* < 0.01, \*\*\**P* < 0.001 and \*\*\*\**P* < 0.0001 compared to DMSO/PBS). **(G-J)** Subchondral bone thickness for each quadrant of the joint measured from µCT images (\**P* < 0.05, \*\**P* < 0.01, \*\*\**P* < 0.001 compared to DMSO/water). **(K)** Representative safranin O-stained sections from each experimental group (Scale bar: 200 µm). Arrows indicate cartilage degeneration of varying severity ranging from fibrillations in the cartilage to loss of surface lamina and vertical clefts/erosion to the calcified cartilage. Data in **B-J** are means ± SD (n = 10 mice/group). *P*-values were determined using one-way Anova with Tukey’s post hoc analysis.

The primary endpoint for this study was to achieve statistically significant reduction in the maximum cartilage degeneration score over the entire joint, in comparison to the DMSO/PBS group (negative control). Only L-006235 gel, but not the equivalent dose of free L-006235 or blank gel showed a statistically significant reduction in the maximum cartilage degeneration score compared to the DMSO/PBS group (Fig. 5B). Histological analysis revealed 43% reduction in the maximum cartilage degeneration score over the entire joint in mice treated with L-006235 gel compared to the DMSO/water treated group (Fig. 5B). Histologically, DMSO/water, free L-006235 and blank gel treated mice showed cartilage degeneration of varying severity ranging from fibrillations in the cartilage to loss of surface lamina and vertical clefts/erosion to the calcified cartilage (Fig. 5K). The L-006235 gel group on the other hand showed minimal or negligible cartilage degeneration (Fig. 5K).

Cartilage degeneration scores in DMSO/water treated mice were higher in the medial quadrants than in the lateral quadrants (Fig. 5, C-F) consistent with a previous report ^47^, which shows that OA in the DMM model is associated with lesions in the central weight bearing areas of the medial femoral condyle (MFC) and medial tibia plateau (MTP). Compared to DMSO/water, L-006235 gel treatment but not the free L-006235 resulted in a significantly lower cartilage degeneration score in the medial quadrants (Fig. 5, C-F). For medial femoral condyle (MFC), blank gel also achieved statistically significant reduction in cartilage degeneration score as compared to the DMSO/PBS group, but the reduction was less pronounced (*P* < 0.05) as compared to the reduction observed with L-006235 gel (*P* < 0.01).

We also measured subchondral bone plate (SBP) thickness, which has been previously shown to increase in OA joints ^49^. SBP thickness was measured for each quadrant of the knee joint on 2D micro-CT grayscale images (Fig. S13) ^49^. Compared to sham surgery, DMM mice treated with DMSO/water showed significantly higher SBP thickness in the medial quadrants but not in the lateral quadrants (Fig. 5G-J), which is consistent with higher cartilage degeneration scores observed in the medial quadrants. Treatment with L-006235 gel but not with free L-006235 significantly reduced the SBP thickness in the medial quadrants, as compared to the DMSO/water group (Fig. 5G-J). Blank gel also showed significant reduction in SBP thickness in the medial quadrants, which is consistent with the moderate reduction in cartilage degeneration score observed in MFC of blank gel treated mice. We hypothesized that the moderate reduction in MFC cartilage degeneration score and reduction in SBP thickness due to blank gel, might be attributable to the lubricating effect of the hydrogel, as demonstrated previously for another fibrous hydrogel^50^. To investigate the lubricating potential of TG-18 hydrogel, we measured the coefficient of friction (µ) between healthy porcine cartilage discs immersed in TG-18 hydrogel or PBS. Using a rotational rheometer and applying shear rates from 0.1 to 1 s^-1^, as per a method described previously^51^ we confirmed a significant lubricating effect of TG-18 hydrogel, evidenced by a 50% reduction in the friction coefficient relative to PBS (Fig. S14B). Moreover, subjecting the cartilage discs immersed in TG-18 hydrogel to pre-conditioning *via* mechanical loading conditions relevant to resting or running human knee joints did not diminish the lubricating effect of the hydrogel (Fig. S14, C-D).

## Discussion

We demonstrated that a TG-18-based mechanically resilient soft hydrogel enables efficient intra-articular delivery of L-006235 in physically active joints, and reduces disease progression in an aggressive PTOA model exacerbated by rigorous physical activity. *In vitro*, L-006235 gel showed rapid self-healing under experimental conditions that simulate continuous human running. Hydrophobic interactions between L-006235 and TG-18, and the rapid dynamic self-healing of the hydrogel were critical to prevent pre-mature drug release during mechanical loading and maintain sustained release afterwards.

TG-18 hydrogel has advantages over previously developed systems for intra-articular DMOAD delivery. Compared to particle-based systems such as liposomes ^15,16^, solid lipid nanoparticles (SLNs) ^52^, micelles ^53^ and polymeric particles ^17–19^, hydrogels in general have the advantage of a cushioning or lubrication effect that helps maintain joint fluidity and minimizes cartilage friction ^54^. We also observed this for the TG-18 hydrogel. Moreover, our approach surpasses methods that utilize tough hydrogels to achieve mechanical stability ^55^, which often require chemical crosslinkers ^56^ that are associated with substantial toxicity concerns ^57^. For example, a mechanically stable hydrogel platform based on a four-arm maleimide-functionalized polyethylene glycol (PEG-4MAL)^55^ requires chemical crosslinking with dithiothreitol (DTT), which can generate reactive oxygen species, leading to cell death and apoptosis ^58,59^. In contrast, TG-18 hydrogel self-assembles through non-covalent, hydrophobic interactions, avoiding the use of toxic chemical crosslinkers. Additionally, TG-18 is commercially available off the shelf at high purity (Good Manufacturing Practice grade) making our approach amenable to large scale manufacturing and readily translatable. TG-18 hydrogel also offers substantial advantages over hyaluronic acid hydrogel – the most commonly reported material for intra-articular delivery of DMOADs ^12–14^. Hyaluronic acid is non-thixotropic and demonstrates poor stability under conditions of mechanical loading in joints ^60,61^, resulting in its rapid clearance within 24-48 h ^62,63^. Therefore, hyaluronic acid often requires chemical modification to improve its mechanical strength and joint residence time ^64^, which may compromise scalability and translatability. Another non-thixotropic hydrogel reported previously for intra-articular drug delivery ^37,38^ and based on gellan gum also showed poor recovery under mechanical loading conditions in our study, resulting in increased drug release afterwards. In contrast, TG-18 hydrogel exhibits dynamic self-healing due to its supramolecular nature, which enables mechanical resilience and prevents the impact of mechanical loading on drug release. Overall, all these qualities highlight TG-18 hydrogel’s superiority for intra-articular drug delivery under conditions of physical activity, and to our knowledge, mark it as the first hydrogel proven to endure human running conditions and enable sustained DMOAD delivery in active joints.

TG-18 hydrogel is not limited to the encapsulation of L-006235. It is a versatile platform that can be easily adapted for the non-covalent encapsulation of other molecules^26^. For this proof-of-concept study, we chose L-006235 as the DMOAD, which prevents cartilage degeneration by inhibiting cathepsin-K – a key enzyme expressed in chondrocytes that mediates PTOA pathology by cleaving type II collagen and aggrecan, the main components of the cartilage matrix ^31–36^. Cathepsin-K inhibitors have shown promise as systemic therapies for a variety of musculoskeletal diseases^33^. However, off-target effects due to high systemic exposure have hindered their clinical development. For example, odanacatib demonstrated excellent therapeutic efficacy in a phase III clinical trial in patients with osteoporosis ^65^. However, an increased risk for adverse cardiovascular events, particularly stroke, terminated further development of this molecule for osteoporosis. In the case of PTOA, we demonstrated local injection of this clinically relevant drug, for instance using the TG-18 hydrogel can be expected to minimize systemic off-target effects and might enable the clinical translation of this promising therapeutic modality. TG-18 hydrogel can also improve the therapeutic efficacy of L-006235 in PTOA, as observed in our study in mice. L-006235 gel showed a 43% and 45% reduction in maximum cartilage degeneration score and subchondral bone thickness, respectively, as compared to DMSO/water. In contrast, free L-006235 did not show a significant reduction in maximum cartilage degeneration score, which is likely due to the rapid clearance of L-006235 from the joints of treadmill running mice. Blank hydrogel also failed to meet the primary endpoint in this study but showed a moderate reduction in cartilage degeneration score compared to DMSO/water in the MFC and reduced SBP thickness in the MFC and MTP compartments. We hypothesized that this is the result of the lubricating properties of the TG-18 hydrogel ^50^, which we confirmed *in vitro*, as evident by >50% reduction in the coefficient of friction between cartilage discs under varied mechanical loading conditions. These additional beneficial properties of TG-18 hydrogel deserve further exploration in the future.

Our study has several strengths. First, to prevent pre-mature release and loss of L-006235 during mechanical loading, we leveraged the non-covalent, hydrophobic interaction of L-006235 with TG-18, instead of chemically conjugating the two molecules. Chemical conjugation would result in a new chemical entity, which may pose regulatory hurdles, and may also negatively impact gelation. Second, we rigorously tested the mechanical properties of TG-18 hydrogel by *in vitro* rheometry. We used strain and frequency parameters relevant to human knees ^41,42^, which enabled us to study hydrogel thixotropy and sustained release of L-006235 under conditions simulating human running. Third, we validated sustained release of encapsulated agents from TG-18 hydrogel *in vivo* using two complementary techniques – IVIS and LC/MS-MS. Fourth, to determine the therapeutic efficacy of L-006235 gel, we used a clinically relevant mouse model of PTOA, with mice subjected to treadmill running. This model mimics physical activity-induced repeated mechanical loading of knee joints that is experienced by patients with PTOA engaging in activities such as running, exercise and playing sports. This model results in aggressive OA progression, as treadmill running of mice at 400 m/30min/day after DMM surgery exacerbates the OA phenotype ^46^. Demonstrating reduction in disease progression in such an aggressive model of PTOA further strengthens the impact of our findings, and strongly conveys the capability of the TG-18 hydrogel for DMOAD delivery. To our knowledge, this is the first report demonstrating DMOAD-mediated reduction of PTOA progression in physically active joints. Finally, we used a Safranin O staining-based scoring system that is recommended by OARSI as the standard method for histological assessments of osteoarthritis in mice ^48^. This scoring system is an established tool to sensitively evaluate OA progression, and enables comparative evaluation. Using the OARSI method, we clearly demonstrated the superior therapeutic effect of L-006235 gel, as compared to free L-006235 and blank gel in terms of reducing the cartilage degeneration score with respect to the DMSO/PBS group (negative control).

Our study also has certain limitations and there are additional questions that need to be answered. First, loading L-006235 into the TG-18 hydrogel at 20 mg/mL concentration resulted in formation of precipitates, which limited the maximum drug loading for this study. We believe that the formation of precipitates is a result of excess drug molecules impacting the self-assembly of TG-18. Higher drug loading might be achievable by tuning the TG-18 concentration, which warrants further investigations in the future. Second, in the rotational rheometer study, we used a single set of strain and frequency values to model human running. Similarly, for the experiments involving treadmill running mice, we only used one running speed and duration, based on a previously published report ^46^. Third, while the rheo-microscope study enabled us to visually confirm the presence of higher order fibrous structures in TG-18 hydrogel under high strain and high frequency conditions, the low resolution of optical microscopy is an inherent limitation of this approach. Imaging techniques that couple rheometry with high resolution microscopy could be used in the future to examine nanoscale changes in the TG-18 hydrogel under conditions of mechanical loading. Fourth, we were unable to compare the TG-18 hydrogel *in vivo* with previously reported hydrogels for intra-articular drug delivery. Mechanical loading in mouse joints is significantly lower compared to human joint, and this discrepancy might prevent significant differences from emerging when comparing our hydrogel platform with previous systems under such conditions. We therefore opted for *in vitro* benchmarking, which allows us to accurately simulate the mechanical loading conditions of human knee joints. Our rigorous experiments mimicking these conditions clearly demonstrated that the advantages of TG-18 hydrogel over a gellan gum-based hydrogel, reported previously for intra-articular applications ^37,38^. For relevant *in vivo* benchmarking in the future studies, larger animal models such as sheep or goats, which have larger joints and mechanical loading conditions similar to those of human joints, are needed. These models would also would permit the injection of larger volumes (2-6 mL instead of 4 µL) more similar to what one would administer in humans ^66^, thus providing a more accurate estimate of joint residence time and therapeutic efficacy in humans. Future efficacy studies should also evaluate the treatment efficacy over longer periods of time (9-12 weeks) and assess clinically relevant late-stage endpoints such as pain alleviation and gait ^68,69^. Lastly, since the mechanism of L-006235 and other cathepsin-K inhibitors in preventing cartilage degeneration in osteoarthritis is well-established ^32–36^, we did not perform any molecular or cellular analyses to interrogate the biological effects of L-006235 gel. The inclusion of experiments to understand the biological mechanism will be imperative in future studies of hydrogels loaded with investigational new drugs.

In conclusion, our study addresses a critical and previously overlooked aspect of intra-articular drug delivery by studying the impact of mechanical loading conditions on the stability of the drug delivery platform and the release kinetics of encapsulated drugs. We have demonstrated that intra-articular delivery of DMOADs using a mechanically resilient TG-18 hydrogel is a promising approach for reducing PTOA progression in physically active joints. Our approach is simple and scalable, and has potential to enable clinical translation of intra-articular therapies based on promising DMOADs for reducing disease progression in physically active patients with PTOA.

## Materials and Methods

### Experimental design

The objective of this study was to test if a thixotropic hydrogel platform with non-covalently encapsulated DMOAD can achieve efficient intra-articular DMOAD delivery to reduce PTOA progression in physically active joints. As a proof-of-concept DMOAD, we used the cathepsin K inhibitor L-006235, which had previously been shown to inhibit OA progression when administered systemically in pre-clinical models of OA ^32^. L-006235 was encapsulated in the TG-18 hydrogel *via* non-covalent, hydrophobic interactions with TG-18 ^32^. We first determined the maximum loading of L-006235 that can be achieved in TG-18 hydrogel without causing drug precipitation. Next, we investigated the *in vitro* release kinetics of L-006235 in synovial fluid derived from OA patients. Using a rotational rheometer, we studied the impact of mechanical loading at levels relevant to running human knees on the viscoelastic properties, drug release and morphology of the L-006235 gel. L-006235 gel was then evaluated for *in vitro* biocompatibility with primary human chondrocytes and fibroblast-like synoviocytes from healthy subjects or OA patients, followed by *in vivo* safety evaluation upon intra-articular administration in mice. We assessed the effect of treadmill running on the release of the fluorescent dye DiR from L-006235/DiR co-loaded hydrogel and compared the pharmacokinetics of L-006235 injected into the mouse knee as free drug or loaded into the TG-18 hydrogel. Finally, we investigated the therapeutic efficacy of L-006235 gel in treadmill running mice induced to develop PTOA by means of DMM surgery. The primary endpoint for the efficacy study was to achieve statistically significant reduction in the maximum cartilage degeneration score over the entire joint, in comparison to the DMSO/PBS group (negative control).

### Hydrogel preparation and drug encapsulation

To prepare 1 mL of blank hydrogel (10% w/v), 100 mg of triglycerol monostearate (TG-18) (AK Scientific) was weighed into a glass scintillation vial, followed by the addition of 1 mL DMSO-water mixture (1:4 volume ratio). The mixture was then heated to 60-80°C with a heat gun until TG-18 dissolved. The vial was placed on a flat surface and allowed to cool for 15-30 min at room temperature, resulting in hydrogel formation. Gelation was complete when no gravitational flow was observed upon inversion of the vial. L-006235 (Tocris Bioscience) without or with 1,1’-dioctadecyl-3,3,3’,3’-tetramethylindotricarbocyanine iodide (DiR) (Thermo Fisher Scientific) was added together with TG-18 to reach a final concentration of 10-20 mg/mL L-006235 (1-2% w/v) or 100 µg/mL DiR (0.01% w/v). To prepare Rhod-B gels, rhodamine-B (Santa Cruz Biotechnology) was added together with TG-18 to reach a final concentration of 15 mg/mL (1.5% w/v). For *in vivo* studies, we transferred the heated hydrogel preparations into microliter glass syringes prior to gelation. The final volume of the hydrogel upon gelation remained same as the initial total volume of TG-18/water/DMSO/drug mixture due to efficient gelation of water molecules between the TG-18 fibers. Hydrogel morphology was characterized by JEOL SEM (JSM-IT500HR)), ASIA PTE. Ltd., Singapore, acceleration voltage, 2kV). To perform electron microscopic analysis, xerogels were prepared by lyophilizing the gels. Small amounts of xerogels were placed on SEM stubs, air dried, and coated with platinum (2 nm thickness) using the Auto Fine Coater (JEC-3000FC) JEOL, ASIA PTE. Ltd., Singapore).

To prepare L-006235-loaded gellan hydrogels, gellan gum (0.4% w/v) was first dissolved in distilled water. In a scintillation vial, 400 µL of gellan gum solution was mixed with 400 µL of L-006235 solution (75 mg/mL) prepared in DMSO-water mixture (1:4 volume ratio). 200 µL of this mixture was then added to 300 ul of simulated synovial fluid, prepared as described previously ^70^, followed by stirring until gelation happened.

### *In vitro* drug release assay

L-006235 gel (50 µL, 15 mg/mL) was placed in dialysis bags (8-10 kDa molecular weight cut-off) (Spectrum Labs) and suspended in PBS (550 µL). 200 µL synovial fluid (100%) from human OA joints (Articular Engineering) or an equal volume of PBS was added to the dialysis bag at multiple time points. Then, the dialysis tubes filled with hydrogel in release medium were placed in 40 mL sink medium (PBS), and incubated at 37°C with a shaking speed of 150 rpm. At each time point, an aliquot (1 mL) of sink medium was removed and replaced with the same volume of fresh PBS to ensure constant sink conditions. Aliquots were lyophilized and dissolved in 250 µL DMSO followed by quantification by high-performance liquid chromatography (HPLC) (Agilent 1260 Infinity II Quaternary LC pump liquid chromatography system, Zorbax SB C-18 column, 250 x 4.6 mm, 5 µm). Release of L-006235 from gellan hydrogel (500 µL, 15 mg L-006235/mL) using the same protocol.

### *In vitro* rheometry experiments

The impact of mechanical loading on the viscoelastic properties of L-006235 gel was evaluated using a rotational rheometer (Discovery HR-2, TA Instruments) with cone-plate geometry (1°, 40 mm diameter) (TA instruments). 500 µl of L-006235 gel (15 mg L-006235/mL) or blank TG-18 hydrogel and PBS was placed on the plate. We first performed strain sweep measurements at 37°C to characterize define the LVE region for blank and L-006235-loaded hydrogel ^40^. Next, a dynamic frequency sweep was performed between 0.1 to 5 Hz, while keeping the strain constant at 0.1%, which was within the LVE region of both blank and L-006235-loaded hydrogel.

To evaluate the impact of repeated mechanical loading on the viscoelastic properties of L-006235 gel, we first subjected the L-006235 gel at 37°C to low strain and low frequency (0.5%, 0.1 Hz; conditions resembling a resting human knee) for 1 minute followed by 30 consecutive cycles of alternating strain and frequency parameters described previously for human knee joints under resting and running conditions ^41,42^. Each cycle involved 1 minute of high strain high frequency (35%, 2.5 Hz; conditions resembling a running human knee) followed by 1 minute of low strain and low frequency (0.5%, 0.1 Hz). G’ and G’’ were measured over 30 cycles, hydrogel (50 µL) was collected and evaluated for *in vitro* drug release in PBS (37°C), as described above. Hydrogel was also imaged using HR-SEM as described above to study potential morphological changes due to mechanical loading. To determine pre-mature release and loss of L-006235 from the hydrogel due to high strain and high frequency, we centrifuged L-006235 gel at 10,000 g for 10 min after 30 cycles of mechanical loading to separate any free drug resulting from pre-mature release. L-006235 remaining in the gel pellet was quantified using HPLC, as described above. Impact of repeated mechanical loading on the viscoelastic properties of L-006235 gellan gel and L-006235 release was also performed using the same protocol, as described above.

To evaluate if L-006235 gel could recover from high strain and frequency applied for longer duration than 1 min, hydrogel in the presence of PBS was subjected at 37°C to a single cycle of alternating strain and frequency, which involved 2 minutes of low strain and low frequency followed by high strain and high frequency for 5, 10 or 15 minutes, and finally low strain and low frequency for 2 minutes. G’ and G’’ were measured during the entire cycle, hydrogel was collected and evaluated for *in vitro* drug release and imaged using HR-SEM as described above. To determine pre-mature release and loss of L-006235 from the hydrogel, we centrifuged L-006235 gel at 10,000 g for 10 min after each cycle of mechanical loading to separate any free drug resulting from pre-mature release. L-006235 remaining in the gel pellet was quantified using HPLC, as described above.

Impact of mechanical loading on Rhod-B gel was determine by subjecting the gel at 37°C to a single cycle of alternating strain and frequency, involving 5 minutes of low strain and low frequency followed by high strain and high frequency for 15 minutes, and finally low strain and low frequency for 5 minutes. G’ and G’’ were measured over 30 cycles. At the end, Rhod-B gel was centrifuged at 10,000 g for 10 min and dye concentration in the pellet was determined using a fluorescence plate reader. We also evaluated rhodamine-B release from a fresh Rhod-B gel in the absence of mechanical loading conditions but otherwise similar conditions of temperature and sink volume.

Impact of mechanical loading on the blank TG-18 hydrogel was determined by subjecting the hydrogel at 37°C to three consecutive cycles of alternating strain and frequency. Each cycle involved 5 minutes of low strain and low frequency followed by 5, 10 or 15 minutes of high strain and high and finally 5 minutes of low strain and low frequency. To elucidate if TG-18 hydrogel’s thixotropic behavior is an inherent property of the TG-18 molecule or a result of self-assembly of TG-18 into a supramolecular hydrogel, we dissolved TG-18 in DMSO (TG-18/DMSO; 10% w/v) and subjected this non-self-assembled TG-18 mixture at 37°C to 1 minute of low strain and low frequency followed by 20 consecutive cycles of alternating strain and frequency. Each cycle involved 1 minute of high strain and high frequency followed by 1 minute of low strain and low frequency.

To image TG-18 hydrogel real time under mechanical loading conditions relevant to running and resting human knee, we subjected the blank hydrogel in the presence of PBS at 37°C to 30 consecutive cycles of alternating strain and frequency parameters using a rheo-microscope (Anton-Paar), which combines light microscopy imaging with rheological measurements. Each cycle involved 15 seconds of low strain and low frequency followed by 10 seconds of high strain and high frequency. For each cycle, six images at 10X magnification were taken under high strain high frequency conditions, and four images were taken under low strain and low frequency conditions.

### Lubrication test

The coefficient of friction was measured using a method described previously (Fig. S14) ^51^. A disc of porcine cartilage (8 mm by 2 mm) was fixed to the bottom of a sandpaper-covered 40 mm petri dish which was fixed to the base plate of a rotational rheometer (TA Instruments Discovery HR-2, New Castle, DE) by cyanoacrylate glue. A complimentary disc of porcine cartilage was fixed to a 20 mm parallel plate covered with sandpaper, and the petri dish on the base plate of the rheometer was filled with either TG-18 hydrogel or PBS. The top plate was lowered until contact between the cartilage discs was achieved. Flow sweep measurements were performed with shear stress ranging from 0.1 to 1 s^-1^. Shear stress and normal stress were recorded and the ratio of shear stress to normal stress was reported as the coefficient of friction. To determine if mechanical loading conditions relevant to resting and running human knee joints impact the lubricating property of the hydrogel, the cartilage discs immersed in TG-18 hydrogel or PBS were pre-conditioned for 1 minute prior to the flow sweep measurement by applying a strain of 0.5% with a frequency of 0.1 Hz (resting human knee joint) or a strain of 35% with a frequency of 2.5 Hz (running human knee joint).

### *In vitro* biocompatibility assay

Primary human chondrocytes and fibroblast-like synoviocytes from healthy and OA donors (Articular Engineering) were cultured in T-75 flasks (VWR) at 37°C and 5% CO_2_ in chondrocyte or synoviocyte growth medium (Articular Engineering), respectively. Media were supplemented with 10% fetal calf serum (Articular Engineering).

Primary human chondrocytes and fibroblast-like synoviocytes were harvested at 80% confluency and seeded (seeding density 10,000 cells/well) in 96-well plates (Corning). After 48 h of culture, growth medium was replaced with PBS or fresh medium without or with L-006235 gel (at a final concentration of 2 nM, 200 nM or 2 µM of L-006235), free L-006235 (equivalent to L-006235-loaded hydrogel), blank hydrogel (equivalent to L-006235-loaded hydrogel) or DMSO/water (equivalent to L-006235-loaded hydrogel). The final concentration of DMSO was 0.0013% v/v or lower, which is safe for cell-based *in vitro* studies ^71^. After 24, 48 and 72 h, cellular metabolic activity was determined using 2, 3-bis-(2-methoxy-4-nitro-5sulfophenyl)-2H-tetrazolium-5-carboxanilide (XTT) cell proliferation assay kit (American Type Culture Collection) and normalized relative to cells treated with only medium to determine percentage metabolic activity.

### Animals

All *in vivo* experiments were performed with 6-10-week-old male C57BL/6J mice (Jackson Laboratory). Group sizes for each experiment were determined based on the minimal number of animals needed to achieve a significant difference of *P* < 0.05 between experimental groups for the primary outcome based on the results in our preliminary studies. We randomly selected mice from the cage to assign them to different experimental groups. Experiments were performed in a specific pathogen-free animal facility at Brigham and Women’s Hospital. Mice were housed under standard 12 h light-12 h dark conditions with ad libitum access to water and chow. All mouse studies were performed according to institutional and National Institutes of Health (NIH) guidelines for humane animal use and in accordance with the Association for Assessment and Accreditation of Laboratory Animal Care (AAALAC). Protocols were approved by the Institutional Animal Care and Use Committee (IACUC) at Brigham and Women’s Hospital.

### *In vivo* biocompatibility study

*In vivo* biocompatibility of L-006235-loaded hydrogel was evaluated in healthy C57BL/6J mice injected intra-articularly either once or repeatedly. In the single dose safety study, mice were injected into both knees with 4 µL PBS, L-006235 gel, blank hydrogel or free L-006235. The DMSO dose injected in 4 µL of L-006235 gel, blank hydrogel and free L-006235 formulations was 220 µg/kg (for a mouse weighing 20 g), which is substantially lower than the LD_50_ of DMSO (10.9 g/kg) reported previously *via* intraperitoneal route ^72^. Also, since DMSO is gelled between the TG-18 molecules, it would slowly release over time as the hydrogel degrades, thereby minimizing the overall concentration of free DMSO in the joint at a given time. The injection volume of 4 µL was chosen based on the normal synovial volume of mouse knee joints, which is 4-5 µL ^73^. L-006235 was loaded into the hydrogel at the maximum possible concentration of 15 mg/mL, resulting in 60 µg of L-006235 injected in 4 µL L-006235 gel. Two weeks later, injected knees were harvested and processed for histology as described previously ^48,74^. Briefly, dissected knees were decalcified in EDTA for 6 days on a shaker, followed by embedding in paraffin. Coronal sections (4 µm thick) were cut from the anterior to posterior compartment at 200 µm intervals. Sections were stained with H&E (Sigma Aldrich) or safranin O (Sigma-Aldrich). In the repeated dose study, knees were injected with 4 µL PBS or L-006235 gel on day 0, week 2 and week 4, and harvested for histology at week 6. Hematoxylin and eosin-stained sections were observed in a blinded manner for inflammation or other gross evidence for toxicity. Safranin O stained sections were also observed in a blinded manner for any detrimental effects on the cartilage.

### *In vivo* drug release assay

To investigate the impact of mechanical loading on the release of encapsulated agents from TG-18 hydrogel, healthy mice were injected into both knees on day 0 with 4 µL hydrogel co-loaded with L-006235 and a fluorescent dye – DiR (60 µg L-006235, 0.4 µg DiR). One group ran on a treadmill at 400 m/30 min/day, 5 days/week ^46^, while the other group did not run. Every other day, mice were anesthetized *via* isoflurane inhalation, and imaged using IVIS.

To investigate the *in vivo* release kinetics of L-006235 from TG-18 hydrogel in treadmill running mice, free L-006235 or L-006235 gel (4 µL, 60 µg L-006235) was injected into the right knee joint of healthy mice on day 0. Mice were pre-conditioned to treadmill running before the experiment. Mice ran on a treadmill at 400 m/30 min/day, 5 days/week and were euthanized at 1 h, day 1, 6 or 14. The running speed of 400 m/30 min/day was based on a previously published report ^46^. The injected knee tissue and serum were collected to analyze levels of L-006235. Knee tissue was spiked with 10 µL internal standard (IS; 100 µg/mL indinavir), flash frozen in liquid nitrogen and homogenized using a Cellcrusher cryogenic tissue pulverizer (Cellcrusher, Ireland). Homogenized tissue was suspended in methanol, centrifuged to remove debris, and L-006235 was quantified in the supernatant. Serum was spiked with IS (10 µg/mL indiniavir) and processed by solid phase extraction. Briefly, Strata-X^®^ polymeric reversed phase columns (Phenomenex®, Torrance, California) were conditioned with 1 mL methanol, followed by 1 mL water. Serum samples (100 µL) were loaded at 2 mL/min and cartridges were washed with 3 mL of 5% methanol in water. Finally, analytes were eluted in 1.5 mL methanol at 2 mL/min. Methanol was evaporated and samples were reconstituted in 100 µL of methanol for L-006235 quantification. L-006235 was quantified by liquid chromatography tandem mass spectrometry (LC-MS/MS; Agilent 6530 quadrupole time-of-flight mass spectrometer with 1290 infinity binary ultra-performance liquid chromatography system; Zorbax SB C-18 column, 250 x 4.6 mm, 5 µm). The mobile phase (0.45 mL/min) consisted of 0.1% formic acid in water (A) and 0.1% formic acid in acetonitrile (B), with a 9 min gradient of 90-10% A and a total runtime of 16 min. Injection volume was set as 10 µL. Quantification was performed by multiple reaction monitoring (MRM) in positive mode to detect transitions of *m/z* 467 → *m/z* 286 for L-006235 and *m/z* 641 → *m/z* 421 for IS.

### Destabilization of medial meniscus (DMM) model of PTOA

Destabilization of the medial meniscus (DMM) surgery was performed according to an established protocol ^47^. Briefly, under anesthesia, the right knee joint was opened along the medial border of the patellar ligament, followed by cutting the medial meniscotibial ligament (MMTL). During sham surgery, the right knee joint was opened but the MMTL was not severed.

### *In vivo* therapeutic efficacy study

DMM or sham surgery was performed on day 0 as described above. Three weeks later, mice with DMM surgery were injected in the operated right knee with L-006235 gel (4 µL, 60 µg L-006235), free L-006235 (4 µL, 60 µg L-006235), blank hydrogel (4 µL) or DMSO/water (20% v/v DMSO, 4 µL), followed by repeat dosing at week 5 and 7. Since these hydrogels disassemble under inflammation, their intra-articular injection too early after the DMM surgery can result in pre-mature drug release due to surgical inflammation. We observed that a wait time of 3 weeks after the surgery for the first injection allows surgical inflammation to subside, thereby preventing pre-mature drug release. From week 3 to week 8, mice from all groups ran on a treadmill at 400 m/30 min/day, 5 days/week. The running speed of 400 m/30 min/day was based on a previously published report ^46^, and was chosen to exacerbate the OA phenotype in mice after DMM surgery, resulting in aggressive disease progression. Mice were pre-conditioned for treadmill running before the surgery, and were euthanized at week 9 and knees were harvested. After micro-CT imaging, knees were decalcified, sectioned, and stained with Safranin O as described above. Sections were scored for cartilage degeneration using the Osteoarthritis Research Society International (OARSI) system in a blinded manner by two different readers ^48^.

### Micro CT imaging

After euthanasia, injected knees were harvested and fixed in 4% (w/v) paraformaldehyde for one day followed by 70% (v/v) ethanol. The samples were scanned using µCT 35, Scanco Medical (µCT V 6.1) scanner with isotropic voxel size 12 μm, X-ray tube voltage 55 kV, intensity 145 μ A, and integration time of 600 ms. Analysis was performed using the built in software. The knee joints were three-dimensionally reconstructed using a global bone mineral threshold of 380 mg HA/cm^3^ and digitally sectioned along the frontal plane. The load bearing regions of the joint were selected as the region of interest. To determine subchondral bone plate thickness, the cortical bone of the medial tibial plateau was contoured to exclude the calcified articular cartilage and any portion that is part of an osteophyte ^49^. After selecting the middle section of the facet knee joint, the subchondral bone thickness was directly measured from the medial and lateral regions of tibial plateau and femoral condyle using the inbuilt software in Scanco Medical (µCT V 6.1).

### Statistical analysis

Statistical analysis and graphing were done with Graphpad Prism. The two-tailed Student’s *t*-test was used to compare two experimental groups and one-way ANOVA with Tukey’s post hoc analysis was used for comparing more than two groups. To determine the statistical significance of differences in fluorescence signal decay in the *in vivo* experiments with L-006235/DiR co-loaded loaded hydrogel, the area under the curve for individual mice was calculated and group means were compared. To compare in vitro drug release from L-006235 gel in PBS versus OA synovial fluid, we used a two-way ANOVA analysis using time and sink type (PBS versus OA synovial fluid) as the two variables. To determine the statistical significance of differences in cartilage degeneration scores and SBP thickness, one-way ANOVA with Tukey’s post hoc analysis was used, with the mean of each group compared to the mean of the DMSO/PBS treatment group (negative control). A value of *P* < 0.05 was considered statistically significant.

## Acknowledgments

Department of Defense award W81XWH-14-1-0229 (JMK)

National Institutes of Health grant R01AR077718 (NJ)

Disease Targeted Innovative Research Grant from the Rheumatology Research Foundation (JMK, JE)

King Abdulaziz City for Science and Technology through the Center of Excellence for Biomedicine (JMK)

Football Players Health Study grant (JMK, JE, NJ). The Football Players Health Study is funded by a grant from the National Football League Players Association. The content is solely the responsibility of the authors and does not necessarily represent the official views of Harvard Medical School, Harvard University or its affiliated academic health care centers, the National Football League Players Association, or the Brigham and Women’s Hospital.

## Author contributions

Conceptualization: NJ, JMK, JY, JE.

Methodology: NJ, JY, JE, MD, KS, YW.

Investigation: NJ, JY, MD, KS, YW, DW, TU, VP, MXC, SK, SB, PS, CK, NES, JNL, TC, JJ, JPE, LB, AAH, HAA, MD, EW, RCSY, and JG.

Supervision: NJ, JE, JMK.

Writing—original draft: NJ, MD, KS.

Writing—review & editing: NJ, JE, JMK, JY.

## Declaration of interests

J.M.K. has been a paid consultant and or equity holder for multiple biotechnology companies (listed here: https://www.karplab.net/team/jeff-karp). The interests of J.M.K. were reviewed and are subject to a management plan overseen by his institutions in accordance with its conflict of interest policies. N.J., J.M.K., K.S., S.M. and N.E.S. have pending and issued patents on the hydrogel platform described in this manuscript. The authors declare that they have no other competing interests.

