## Supplementary file for "A Mechanically Resilient Soft Hydrogel Improves Drug Delivery for Treating Post-Traumatic Osteoarthritis in Physically Active Joints"

**This PDF file includes:**

Figs. S1 to S14
Table S1

**Other Supplementary Material for this manuscript includes the following:**

Movie S1

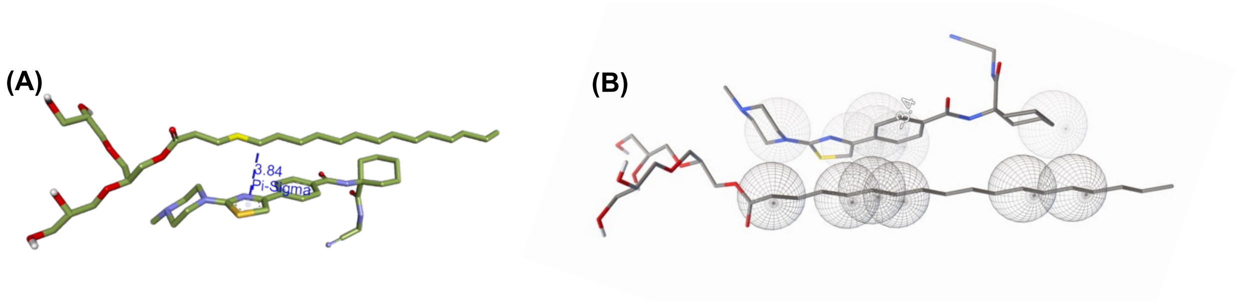


**Fig. S1. Molecular docking studies showed possible hydrophobic molecular interactions between TG-18 and L-006235. (A)** Pi-Sigma hydrophobic molecular interactions were observed with an intermolecular distance of 3.84 Å using Discovery studio visualizer (DSV). **(B)** Vander Waal interactions between TG18 and L006235 as predicted by Autodock Tools (ADT).

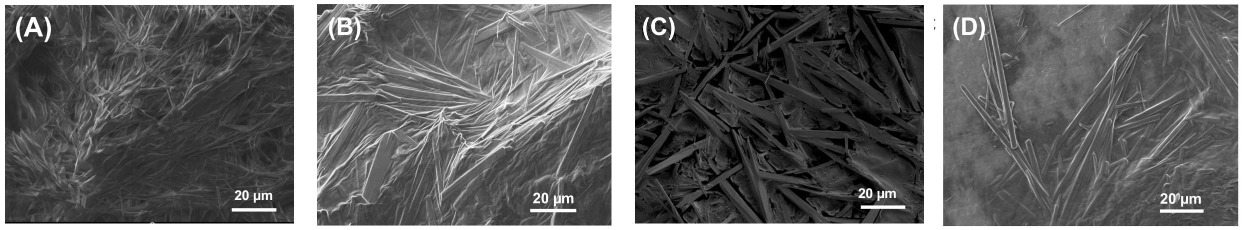


**Fig. S2. TG-18 hydrogel demonstrates fibrous assembly up to 15 mg/ml of L-006235.** High-resolution scanning electron microscopy (HR-SEM) of **(A)** Blank TG-18 hydrogel, and TG-18 hydrogel loaded with **(B)** 10 mg/ml L-006235, **(C)** 15 mg/ml L-006235 and **(D)** 20 mg/ml L-006235.

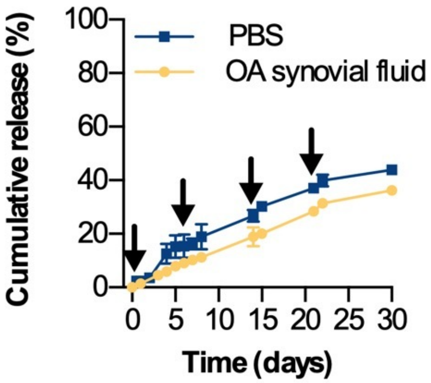


**Fig. S3. L-006235 gel exhibits sustained drug release.** *In vitro* release of L-006235 from L-006235 gel in PBS or synovial fluid from human osteoarthritis (OA) joints. 200 µl of PBS or fresh synovial fluid from human OA knees (100% concentration) were added at the indicated time points (arrows). The difference between release kinetics in PBS vs. OA synovial fluid was not statistically significant over the entire time course ($P$ = 0.1487). Data are means $\pm$ SD of technical repeats (n = 3, each experiment performed at least twice). $P$-value was determined using two-way Anova with Bonferroni correction.

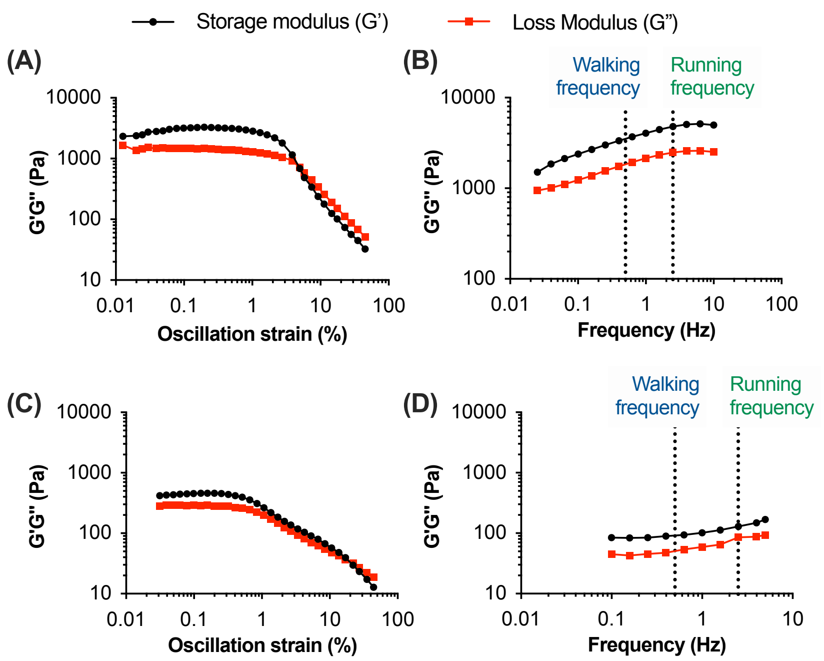


**Fig. S4. L-006235-loaded hydrogel has linear viscoelastic (LVE) region up to 5% strain.** Storage modulus (G') and loss modulus (G") measured during the strain sweep of (**A)** DMOAD-loaded hydrogel and **(C)** blank hydrogel in PBS, performed with a fixed frequency of 0.1 Hz. Storage modulus (G') and loss modulus (G") measured during the dynamic frequency sweep of **(B)** DMOAD-loaded hydrogel in PBS or **(D)** Blank hydrogel in PBS measured at 0.1% strain. Data in **(A-D)** are from a single experiment (repeated three times).

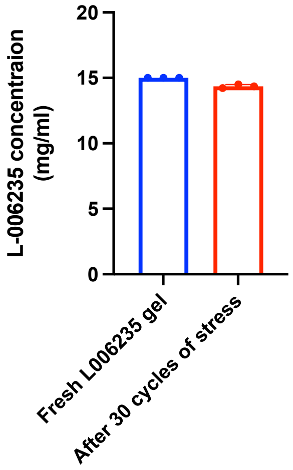


**Fig. S5. Concentration of L-006235 in fresh L-006235 gel or L-006235 gel subjected to 30 consecutive cycles of alternating strain and frequency.** Using a rotational rheometer, L-006235 gel in the presence of PBS was first subjected at 37°C to low strain and low frequency (0.5%, 0.1 Hz; conditions resembling a resting human knee) for 1 minute followed by 30 consecutive cycles of alternating strain and frequency. Each cycle involved 1 minute of high strain and high frequency (35%, 2.5 Hz; conditions resembling a running human knee) followed by 1 minute of low strain and low frequency (0.5%, 0.1 Hz). After 30 cycles, gel was collected, and centrifuged to separate drug released during mechanical loading. L-006235 remaining in the gel pellet was quantified using HPLC.

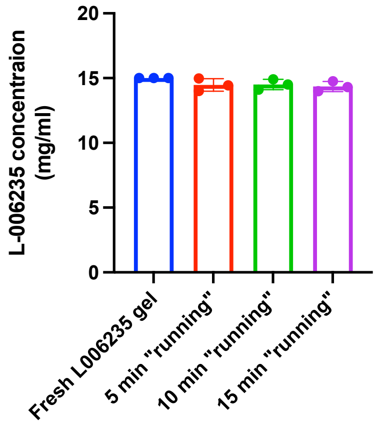


**Fig. S6. Concentration of L-006235 in fresh L-006235 gel or L-006235 gel subjected to a single cycle, with 5, 10 or 15 min step of high strain and high frequency.** Using a rotational rheometer, L-006235 gel in the presence of PBS was subjected to a single cycle of alternating strain and frequency, involving 2 minutes of low strain and low frequency followed by high strain and high frequency for 5, 10 or 15 minutes, and finally low strain and low frequency for 2 minutes. After each cycle, gel was collected, and centrifuged to separate drug released during mechanical loading. L-006235 remaining in the gel pellet was quantified using HPLC.

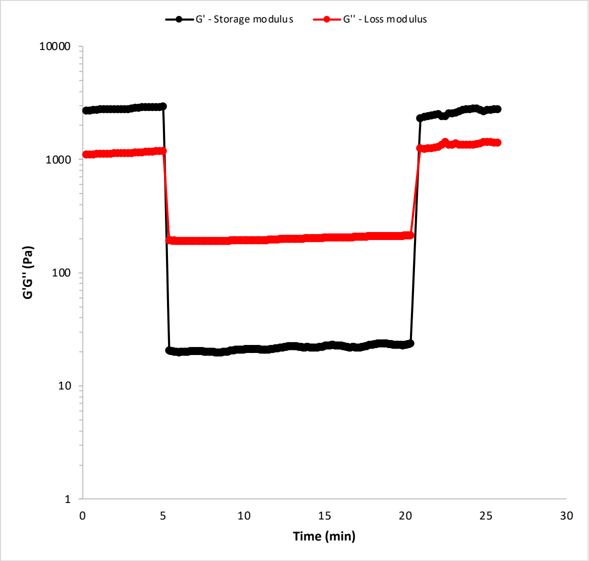


**Fig. S7. Rhodamine-B loaded hydrogel (Rhod-B gel) recovers rapidly after exposure to mechanical loading at levels relevant to running human knee joints.** Rhod-B gel in the presence of PBS was subjected to a single cycle of alternating strain and frequency, involving 5 minutes of low strain (0.5%) and low frequency (0.1 Hz) (conditions resembling a resting human knee) followed by 15 minutes of high strain (35 %) and high frequency (2.5 Hz) (conditions resembling a running human knee) and finally 5 minutes of low strain (0.5%) and low frequency (0.1 Hz). Shear storage modulus (G') and shear loss modulus (G") were measured.

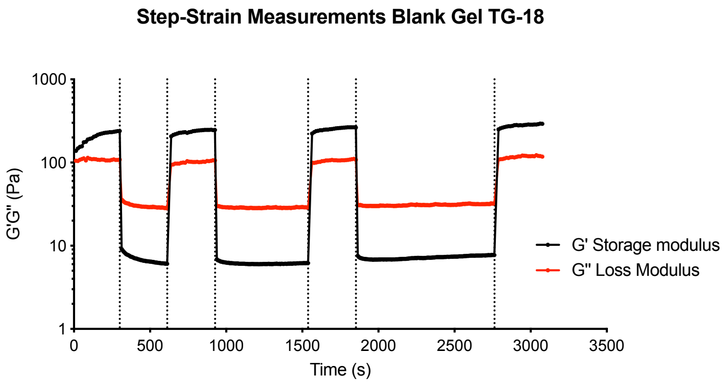


**Fig. S8. Blank TG-18 hydrogel recovers rapidly after exposure to mechanical loading at levels relevant to running human knee joints.** Blank TG-18 hydrogel in the presence of PBS was subjected to three consecutive cycles of alternating strain and frequency. Each cycle involved 1 minute of low strain (0.5%) and low frequency (0.1 Hz) (conditions resembling a resting human knee) followed by 5, 10 or 15 minutes of high strain (35 %) and high frequency (2.5 Hz) (conditions resembling a running human knee) and finally 1 minute of low strain (0.5%) and low frequency (0.1 Hz). Shear storage modulus (G') and shear loss modulus (G") were measured.

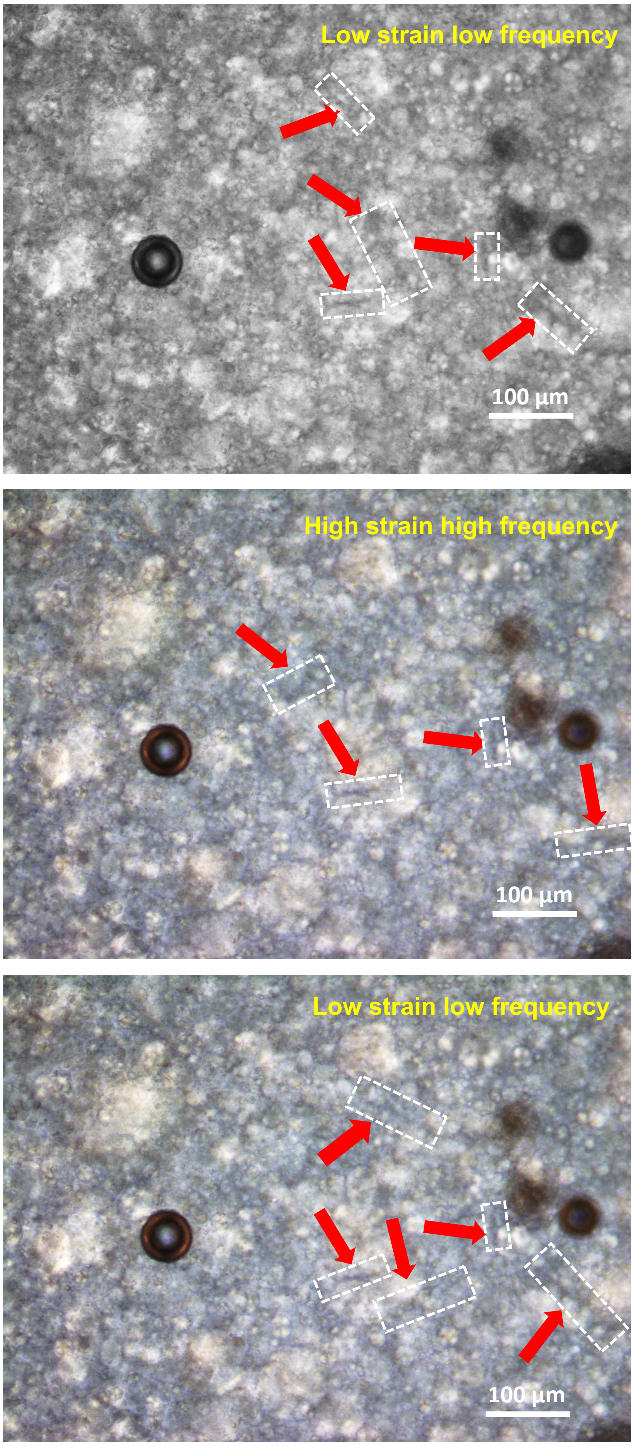


**Fig. S9. TG-18 hydrogel maintain higher order fibrous (thread like) structures under conditions of mechanical loading  relevant to running human knee joints.** Using a rheo-microscope, blank TG-18 hydrogel in the presence of PBS was subjected at 37$^{o}$C to 30 consecutive cycles of alternating strain and frequency parameters. Each cycle involved 15 seconds of low strain and low frequency (0.5%, 0.1 Hz ; conditions resembling a resting human knee) followed by 10 seconds of high strain and high frequency (35%, 2.5 Hz; conditions resembling a running human knee). For each cycle, six images at 10X magnification were taken under high strain and high frequency conditions, and four images were taken under low strain and low frequency conditions. Representative images of blank hydrogel are shown for one cycle. Fibrous structures were observed under both low strain and low frequency, and high strain and high frequency conditions. Entangled fibers are seen as black-colored cob-web like structures on the white background. Prominent fibrous structures are marked with dashed boxes and arrows.

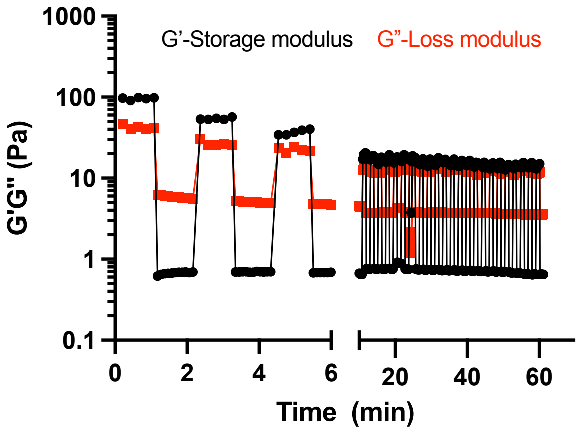


**Fig. S10. Viscoelastic properties of TG-18 dissolved in DMSO (TG-18/DMSO) do not recover upon removing high strain and high frequency.** Using a rotational rheometer, TG-18/DMSO in the presence of PBS was first subjected at 37°C to 1 minute of low strain and low frequency (0.5%, 0.1 Hz; conditions resembling a resting human knee) followed by 20 consecutive cycles of alternating strain and frequency. Each cycle involved 1 minute of high strain and high frequency (35%, 2.5 Hz; conditions resembling a running human knee) followed by 1 minute of low strain and low frequency (0.5%, 0.1 Hz). Shear storage modulus (G') and shear loss modulus (G") were measured.

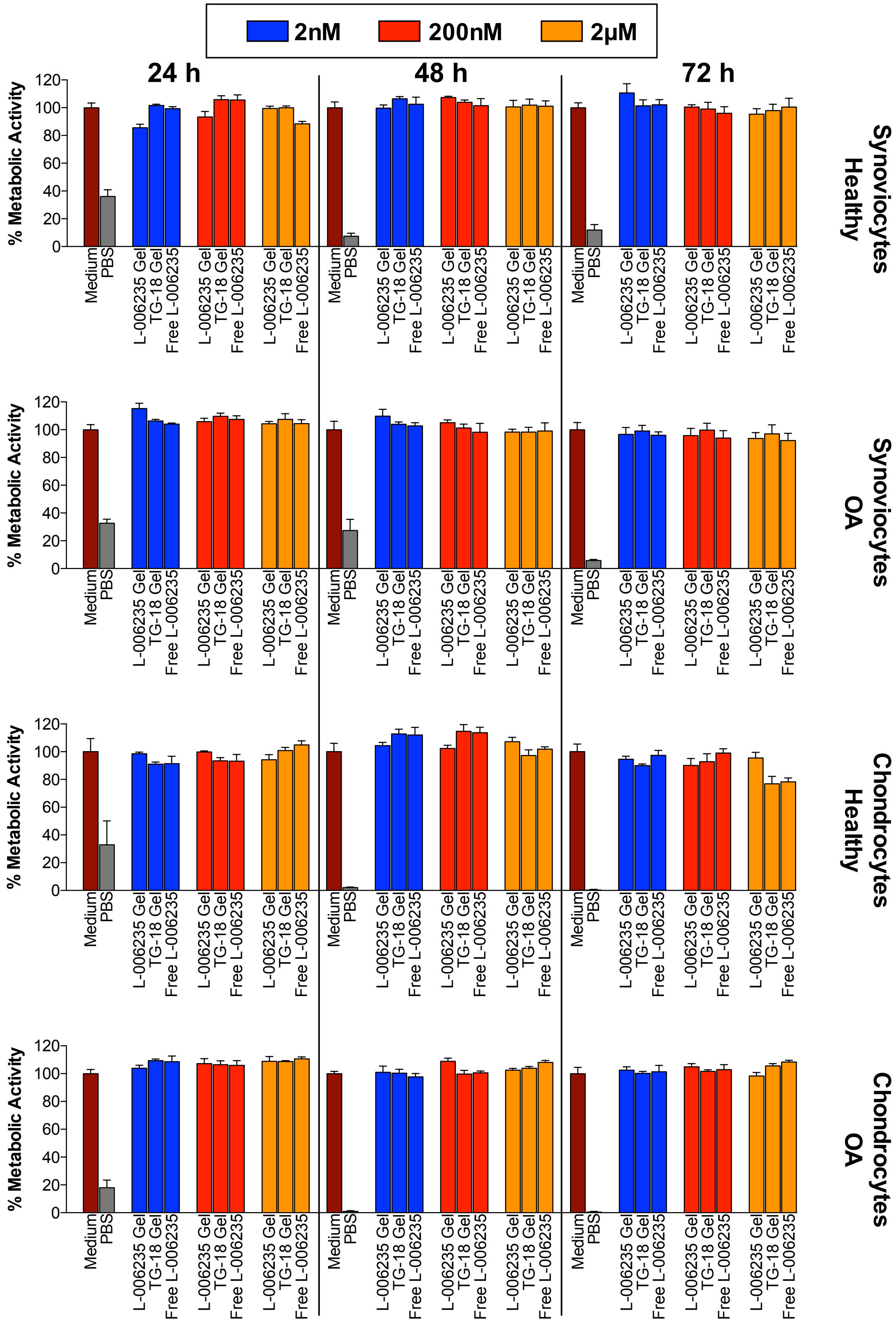


**Fig. S11. L-006235-loaded hydrogel is biocompatible with cells from human joints.** Primary human chondrocytes and synoviocytes from OA or healthy joints were incubated in a 96-well plate in medium, PBS or in medium with L-006235-loaded hydrogel (L-006235 gel) or free L-006235 at 2 nM, 200 nM or 2 μM final concentration of L-006235. Cells incubated in medium with blank TG-18 hydrogel (Blank gel) at a volume equivalent to L-006235 gel were evaluated as controls. After 24, 48 or 72 h of incubation, cellular metabolic activity was measured using the XTT assay. Data in **A-D** are means ± SD of technical repeats (n = 8 wells per condition, experiment performed twice).

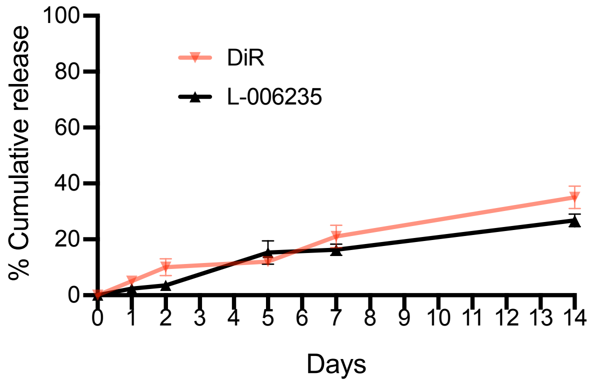


**Fig S12. L-006235 and DiR exhibit similar release kinetics from the TG-18 hydrogel.** *In vitro* release of L-006235 and DiR from L-006235 gel and L-006235/DiR co-loaded TG-18 hydrogel, respectively in PBS. Data are means ± SD of technical repeats (n = 3, each experiment performed at least twice).

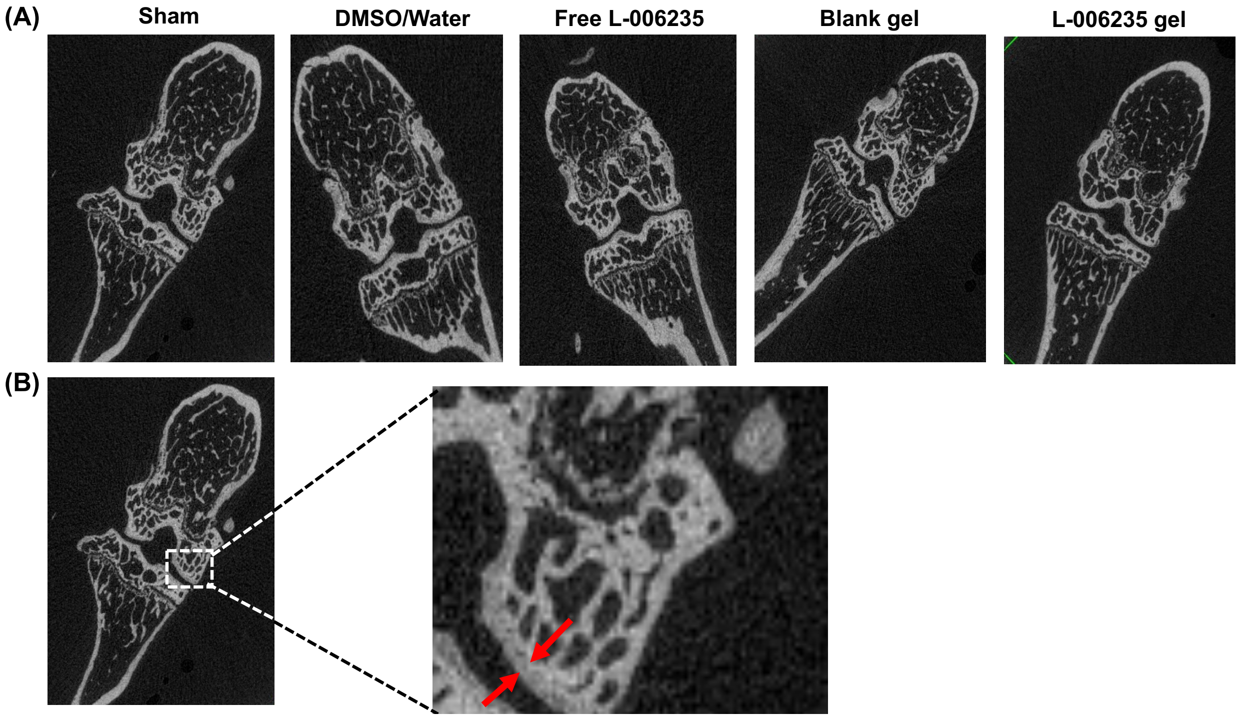


**Fig. S13: (A) Representative μCT scan (2D) of a knee joint for each group.** To determine subchondral bone plate thickness, the cortical bone of the medial and lateral tibial plateau, medial and lateral femoral condyle was contoured excluding the calcified articular cartilage and any portion that is part of an osteophyte. After selecting the middle section of each contoured region, the subchondral bone thickness was measured using the inbuilt software in Scanco Medical (μCT V 6.1). **(B)** Zoomed in image showing medial femoral condyle of a knee joint from sham group, with arrows marking the bone plate thickness.

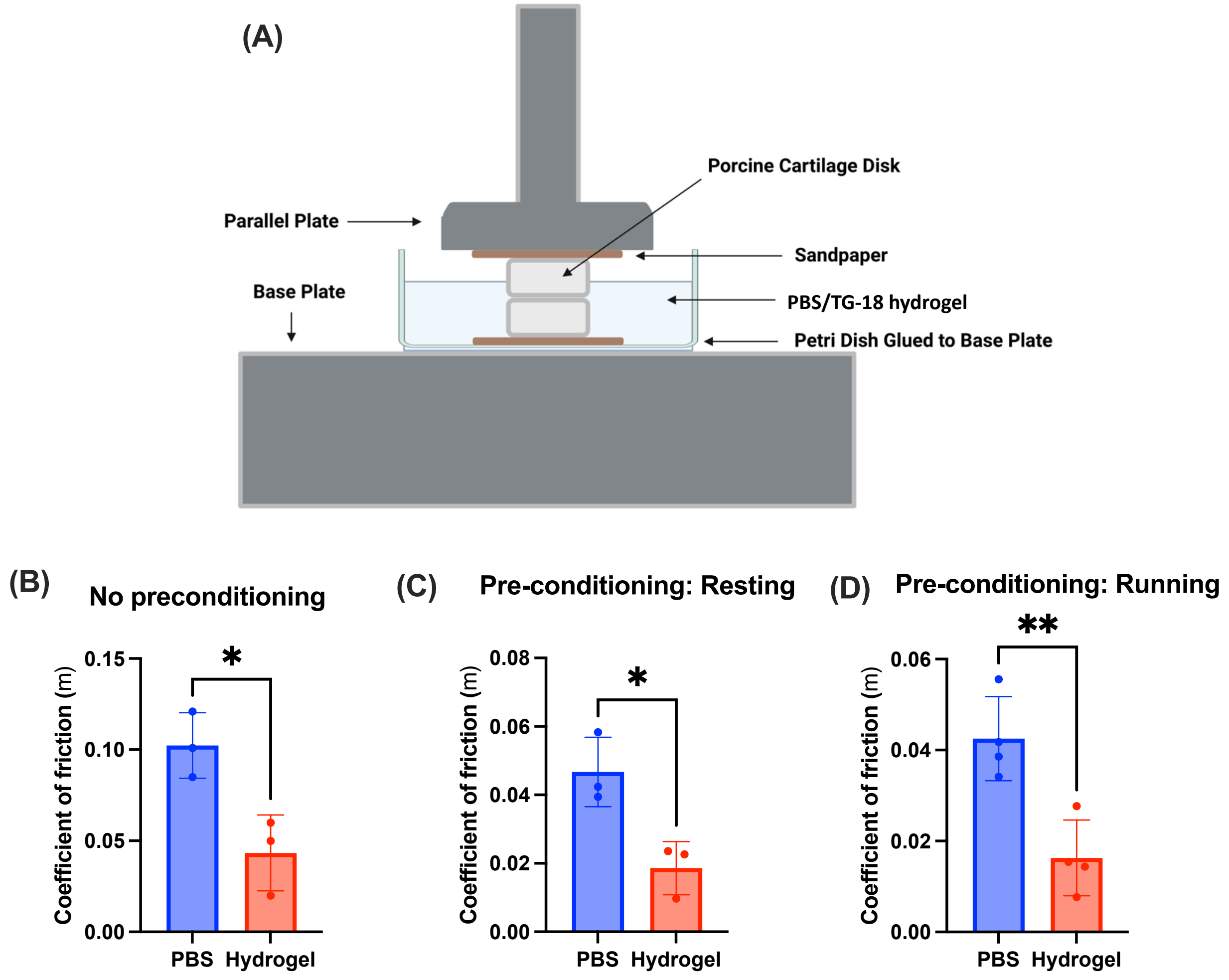


**Fig. S14: TG-18 hydrogel has a lubricating effect and reduces the coefficient of friction between cartilage discs. (A)** Schematic of the set-up used to measure the friction coefficient between two discs of healthy porcine cartilage immersed in TG-18 hydrogel or PBS. The coefficient of friction between the cartilage discs immersed in TG-18 hydrogel or PBS was measured at a shear rate of 0.1-1 $s^{-1}$ **(B)** without any pre-conditioning (*$P < 0.05$), **(C)** after pre-conditioning with mechanical loading mimicking resting human knee joints (*$P < 0.05$) or **(D)** after pre-conditioning with mechanical loading mimicking running human knee joints (**$P < 0.01$). Data in **(B)**, **(C)** and **(D)** are means $\pm$ SD of technical repeats (n = 3-4, each experiment performed at least twice). *P* values were determined using Student's *t*-test.

**Table 1:** Binding affinities of the nine possible conformers of TG-18 and L-006235.

| Conformer | Binding affinity $\Delta G$ (kCal/mol) | Distance from best conformer | |
|---|---|---|---|
| | | rmsd l.b. | rmsd u.b. |
| 1 | -3.4 | 0.000 | 0.000 |
| 2 | -3.3 | 3.592 | 4.561 |
| 3 | -3.2 | 1.981 | 2.310 |
| 4 | -3.2 | 2.411 | 3.053 |
| 5 | -3.1 | 1.649 | 1.903 |
| 6 | -3.1 | 4.445 | 6.375 |
| 7 | -3.1 | 5.012 | 6.608 |
| 8 | -3.0 | 2.438 | 3.071 |
| 9 | -3.0 | 2.558 | 3.263 |

**Movie S1:** Injectability of L-006235 gel through a 27 gauge needle.
